# CMTR1 is recruited to transcription start sites and promotes ribosomal protein and histone gene expression in embryonic stem cells

**DOI:** 10.1101/2022.02.01.478435

**Authors:** Shang Liang, Joana Clara Silva, Olga Suska, Radoslaw Lukoszek, Rajaei Almohammed, Victoria H Cowling

## Abstract

CMTR1 (cap methyltransferase 1) catalyses methylation of the first transcribed nucleotide of RNAPII transcripts (N1 2’-O-Me), creating part of the mammalian RNA cap structure. In addition to marking RNA as self, N1 2’-O-Me has ill-defined roles in RNA expression and translation. Here we investigated the gene specificity of CMTR1 and its impact on RNA expression in embryonic stem cells. Using chromatin immunoprecipitation, CMTR1 was found to bind to transcription start sites (TSS) correlating with RNAPII levels, predominantly binding at histone genes and ribosomal protein (RP) genes. Repression of CMTR1 expression resulted in repression of RNAPII binding at the TSS and repression of RNA expression, particularly of histone and RP genes. In correlation with regulation of histones and RP genes, CMTR1 repression resulted in repression of translation and induction of DNA replication stress and damage. Indicating a direct role for CMTR1 in transcription, addition of recombinant CMTR1 to purified nuclei increased transcription of the histone and RP genes. CMTR1 was found to be upregulated during neural differentiation and there was an enhanced requirement for CMTR1 for gene expression and proliferation during this process. We highlight the distinct roles of the cap methyltransferases RNMT and CMTR1 in target gene expression and differentiation.

## INTRODUCTION

The RNA cap is added to the initiating nucleotide of RNA pol II (RNAPII) transcripts. It protects RNA from degradation by blocking exonucleases during transcription and identifies RNA as cellular in origin or “self”, protecting it from degradation by proteins of the innate immunity pathways (1–3). The most prevalent cap structure, denoted ^m7^GpppX_m_, includes 7-methylguanosine (^m7^G) and first nucleotide ribose O2-methylation (Xm or N1 2’-O-Me) (4,5). N1 2’-O-Me prevents RNA interactions with proteins of the innate immune pathways, e.g. IFIT proteins and RIG-I, which bind to the cap in the absence of N1 2’-O-Me and direct RNA degradation (3,6,7). Proteins involved in splicing, nuclear export, RNA processing and translation initiation are also recruited to RNA via the cap (8,9). The role of N1 2’-O-Me in these gene expression processes is unclear; it may influence the recruitment of cap binding proteins which mediate processing events, including translation, although mechanistic information is lacking (10,11). Recent findings demonstrated superior translation of capped RNA with N1 2’-O-Me in certain cell types, although this did not correlate with binding of the translation initiation factor eIF4E, indicating the influence of other factors (12).

Following transcription initiation, a series of enzymes catalyse RNA cap formation, starting with guanosine linkage to the first transcribed nucleotide by a 5’ to 5’ triphosphate bridge (1,8). This cap guanosine is methylated at the N-7 position, the first and second transcribed nucleotides are methylated on the ribose at the O-2 position, and if the first nucleotide is adenosine it may also be methylated at the N-6 position (13). In mammals, cap guanosine N-7 methylation and first transcribed nucleotide ribose O-2 methylation are catalysed by the cap methyltransferases RNMT and CMTR1, respectively, to form the predominant cap structure, ^m7^GpppX_m_ (4,5).

The N1 2’-O-Me methyltransferase, CMTR1 (ISG95, FTSJD2, KIAA0082)(14,15), was first identified in screens for interferon stimulated genes and later was characterised as a cap methyltransferase (16–19). As an interferon stimulated gene (ISG), CMTR1 has been found to be important for the expression of a subset of other ISGs (20). In vivo, CMTR1 is required for neuromorphogenesis and brain development, in particular promoting Camk2a expression (21). The mechanisms directing these gene-specific impacts of CMTR1 are not known. CMTR1 has a central methyltransferase core surrounded by several protein:protein interaction domains (22,23). It is recruited directly to RNAPII C-terminal domain (CTD) via an interaction with the WW domain and this interaction increases when the CTD is phosphorylated on serine-5 of the heptad repeats (22,23). Approximately half of the CMTR1 in the nucleus is present in a complex with the helicase DHX15 which influences RNA translation (23,24). DHX15 represses the catalytic activity of CMTR1 but does bind to RNAPII or influence the RNAPII-CMTR1 interaction.

In order to investigate the role of CMTR1 in gene expression, ChIP-seq analysis was performed in murine embryonic stem cells (ESCs). The highest levels of CMTR1 were found bound to the ribosomal protein (RP) genes and histone genes. For these genes, CMTR1 binding corelated with phosphorylation of Serine-5 of RNAPII CTD at the transcription start site (TSS). Repression of CMTR1 resulted in loss of RNAPII at the TSS and repression of ribosomal protein and histone gene expression. Conversely, the addition of recombinant CMTR1 to isolated nuclei resulted in increased transcription of histone and RP genes. In correlation with regulation of histones and RP genes, repression of CMTR1 resulted in induction of markers of DNA replication stress and damage, and repression of RNA translation. There was enhanced dependency on CMTR1 for gene expression and cell proliferation when ESC were induced to differentiate.

## MATERIALS AND METHODS

### Cell culture and differentiation

Murine ESCs (Embryonic Stem cells) were cultured in Glasgow’s Minimum Essential Medium (Life Technologies),10% Knockout Serum Replacement (Invitrogen), 1% Non-Essential Amino Acids (Life Technologies), 2 mM L-Glutamine (Life Technologies), 1 mM sodium pyruvate (Life Technologies), 0.35 μM β-mercaptoethanol (Sigma), 100 U/ml recombinant LIF (DSTT, Division of Signal Transduction Therapies, University of Dundee). ESCs were cultured in 0.1% gelatin-coated flasks at 37°C / 5 % CO_2_. For LIF withdrawal-induced differentiation, 1×10^6^cells were seeded on 0.1% gelatine-coated 10 cm plates in ESC medium minus LIF with medium replacement every two days. For neural differentiation, 1-1.6×10^6^ cells were seeded on 0.1% gelatine-coated 10 cm plates in N2B27 medium composed of 50% Dulbecco’s Modified Eagle Medium/Nutrient Mixture F-12 (DMEM/F12) / 50% Neurobasal Medium (Life Technologies), supplemented with 1% B-27 supplement (Life Technologies), 0.5% N-2 supplement (Life Technologies), 2 mM L-Glutamine (Life Technologies) and 0.35 μM β-Mercaptoethanol (Sigma) (25). Medium was replaced every day.

### CRISPR/Cas9 genome targeting

Paired guide RNAs (gRNAs) targeting the end of *Rnmt* and *Cmtr1* gene exon 2 were designed (Figure S1). List of gRNAs used for editing:

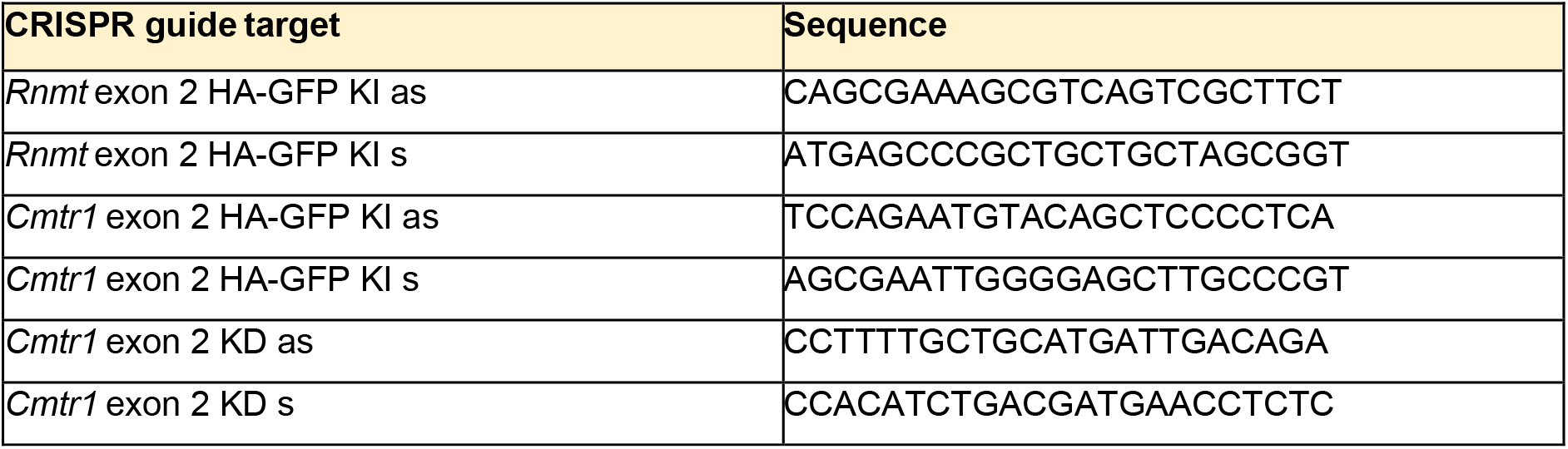

Antisense gRNAs were cloned into pX335-Cas9-D10A. Sense gRNAs were cloned into pBABED-Puro-U6. ESC cells at ~70% confluency were transfected using Lipofectamine 2000 (Life Technologies) and 1μg gRNAs vectors, 3 μg donors. Cells were selected using 1.5 μg/mL puromycin for 2 days. HA-GFP-Rnmt/Cmtr1 KI cells were sorted and GFP positive cells retained. Cells were plated at reduced density in medium supplied with 20% pre-conditioned medium. Single cell colonies were expanded and protein and DNA analysed. For HA-GFP KI ESCs, clones showing no wild-type RNMT/CMTR1 band, but with product approximately 30 kDa larger than the wild-type protein were selected for the genotyping. For *Cmtr1^KD^* ESCs, clones exhibiting significant CMTR1 protein reduction were selected for genotyping. 200 ng genomic DNA was used in genotyping. Primers were designed to ensure correct insertion of HA-GFP encoding cassette for HA-GFP KI ESCs and knockout of *Cmtr1* for *Cmtr1^KD^*ESCs. For HA-GFP-CMTR1 KI ESCs, one *Cmtr1* allele has the designed mutation to create the fusion protein and the other allele has a deletion which prevents expression. Similarly the Cmtr1^KD^ cells have one allele with the KD mutation and one allele with a deletion which prevents expression. To generate ESCs stably expressing HA-CMTR1 in the *Cmtr1^KD^* background, 1×10^6^ *Cmtr1^KD^* cells were transfected with 2 μg pPyPCAGGS vector (empty-vector or HA-CMTR1) using Lipofectamine 2000 (Life Technologies). Successfully transfected cells were selected with puromycin (1 μg/mL).

#### siRNA transfection

siRNA oligos were from Dharmacon, siGENOME Mouse collection. Transfections performed with 50 μM siRNAs using Lipofectamine RNAiMAX (Life Technologies). For RNA-seq /protein analysis, siRNA transfection mix was added to 3 - 4 x10^5^ freshly seeded cells on a 6-well plate. For nuclear run-on and ChIP-seq, siRNA transfection mix was added to 3 - 4 x10^6^ freshly seeded cells on a 10 cm plate. Cells were cultured for 36 hrs (medium replaced after 12 hrs).

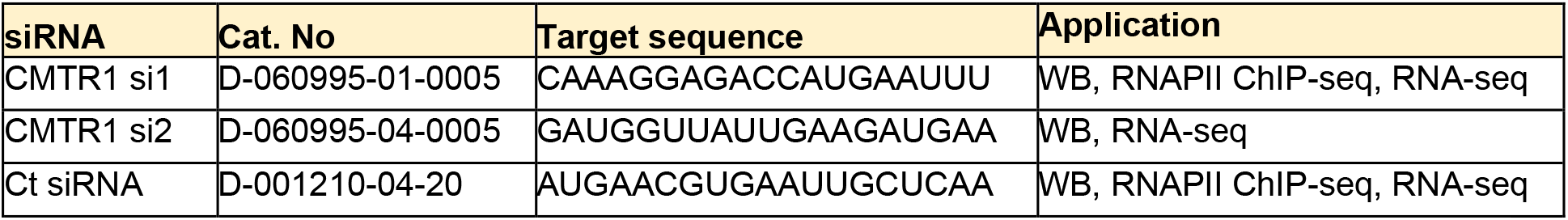

### Proteosome inhibition with MG132

2 – 3 x10^5^ cells were seeded on 6-well plates one day before the treatment. Culture medium was replaced with ESC medium supplemented with 5 μM of MG132 (Calbiochem) or mESC medium with DMSO (as controls). MG132-treated cells were collected after 2, 4 and 6 hrs of treatment.

#### Cell lysate preparation and western blot

2 - 3 x10^6^ cells were seeded on 10 cm plates two days before lysis. For protein analysis, cell lysates were prepared in 10 mM Tris-HCl pH 7.0, 50 mM NaCl, 30 mM sodium pyrophosphate, 50 mM NaF, 5 μM ZnCl_2_, 10% glycerol, 0.5% Triton X-100, 1 mM DTT, 10 mM leupeptin, 1 mM pepstatin, 0.1 mg/mL aprotinin, and phosphatase inhibitor 2 and 3 cocktail (Sigma) as previously described (26).1 μg/μl protein lysates in Laemmli sample buffer supplemented with 0.025 M DTT were boiled for 5 min. 6 −10 μg protein was resolved on a polyacrylamide gel. PVDF (Millipore) was used for electroblotting. Membranes were blocked for 1 hr before overnight incubation with primary antibodies at 4°C. Membranes were washed and HRP-conjugated secondary antibodies (Thermo Fisher) added for 1 hr before imaging using SuperSignal West Pico Chemiluminescent Substrate (Thermo Fisher).

| Antibody antigen | Type | Supplier/ Cat. no |
| --- | --- | --- |
| RNMT | Sheep polyclonal | DSTT |
| CMTR1 | Rabbit polyclonal | Sigma HPA029979 |
| RAM | Sheep polyclonal | DSTT |
| Sox2 | Mouse monoclonal | CST 4900 |
| Oct4 | Rabbit polyclonal | Abcam ab18976 |
| Nanog | Rabbit polyclonal | Abcam ab80892 |
| Klf4 | Goat polyclonal | R&D AF3158 |
| Sox1 | Rabbit polyclonal | CST 4194 |
| DNMT3B | Rabbit polyclonal | Abcam ab122932 |
| GFP | Mouse monoclonal | Roche 11814460001 |
| HA | Mouse monoclonal | BioLegend 901501 |
| RNAPII total (Rpb1) | Rabbit monoclonal | CST 14958 |
| RNAPII S5P | Rat monoclonal | Chromotek 3E8 |
| RNAPII S2P | Rat monoclonal | Chromotek 3E10 |
| Actin | Mouse monoclonal | Abcam ab3280 |
| DHX15 | Rabbit polyclonal | Abcam ab254591 |
| UBC | Rabbit polyclonal | Dako Z0458 |
| Chk1 | Mouse monoclonal | Santa Cruz sc-56291 |
| p-Chk1 | Rabbit monoclonal | CST 2348 |
| γH2AX | Rabbit polyclonal | CST 2577 |

#### Immunoprecipitation

For immunoprecipitation using endogenous antibody, cell lysates were prepared as previously described (26). 2 μg of anti-CMTR1 (Sigma HPA029979) or anti-RNMT (DSTT) antibody or IgG control (DSTT) were added to 1-2 mg of lysates for overnight incubation at 4 °C before the incubation with protein G Dynabeads (Life Technologies) for additional 2 hrs. Beads were washed and eluted as described (25). For HA immunoprecipitation, cells were lysed in PBS, 1 M NaCl, 0.5% v/v NP-40 as described (27). 20 μL anti-HA magnetic beads (Pierce) were added to lysates for overnight incubation at 4 °C. Beads were washed and eluted in TE/1% SDS for 30 min with rotation at room temperature (27).

#### Immunofluorescent staining

Immunofluorescence staining was performed using polyclonal rabbit anti-GFP (1/250, MBL International 598), polyclonal rabbit anti-RNMT (1/250, DSTT), and polyclonal sheep anti-CMTR1 (1/50, DSTT) antibodies. Cells plated on Ibidi 8-well chamber slides were fixed with 4% paraformaldehyde for 20 min, permeabilised with 1% Triton/PBS for 5 min and blocked with 5% donkey serum/PBS for 1 h at room temperature. Primary antibodies diluted in either blocking buffer or 2.5% BSA/PBS were incubated with cells overnight at 4°C. Immunofluorescence labelling was performed using appropriate secondary antibodies conjugated to Alexa fluor 594 dyes (Life Technologies). Stained cells were mounted in Ibidi mounting medium with DAPI prior to imaging with a Leica SP8 confocal microscope.

#### Annexin V staining

10^6^ cells were seeded on 10 cm plates in mESC medium or mESC medium without LIF. Every 24 hrs, cells were trypsinised, washed and resuspended in Fixable Viability Dye eFluor 780 (eBioscience, Life Technologies) for 10 min on ice. Cells were stained with 5 μL of APC-Annexin V (Biolegend) in 45 μL of 1x Annexin V binding buffer (10 mM Hepes (pH 7.4), 140 mM NaCl, 2.5 mM CaCl_2_) for 10 min at room temperature, protected from light. Cells were subject to flow cytometry with a BD LSR Fortessa Cell Analyser System and analysed using FlowJo software.

#### Alkaline phosphatase staining

Cells were seeded on 6-well plates at a concentration of 70 cells/cm^2^. After 3 days, cells were washed in PBS and medium was replaced with 10-fold decreasing concentrations of LIF (from 100 units/mL to 0 units/mL). After 6 days, cells were fixed and stained with Alkaline Phosphatase Staining Kit (Sigma). Colonies were counted under a Zeiss Axiovert 40C microscope and scored into three distinct categories: stained (pluripotent), mixed and unstained (differentiated).

#### ^35^S cysteine and methionine cell labelling

60-70% confluent cells from 6-well plates were used for the labelling. Cells were incubated with mESC medium supplemented with 0.8 Mbq/mL of ^35^S Protein Labeling Mix (Perkin Elmer) at 37 °C with 5 % CO_2_ for 30 min. Cells were lysed, lysates were loaded into SDS-PAGE gels as described in the previous section. Then the gel was fixed with 30% methanol and 7% acetic acid for 40 min. The fixed gel was dried using a vacuum gel dryer for 1 h. Then the dried gel was exposed to a phosphor plate before the scanning using Typhoon imager (GE).

#### Detection of mRNA cap structures by Mass Spectrometry

Performed according to (4).

#### ChIP

2 - 3 x10^6^ cells were seeded on a 10 cm plate one or two day(s) before the ChIP experiment. Cells were cross-linked with 1% formaldehyde for 10 min. Glycine was added to quench the reaction. The cell lysates were prepared as described (27). Chromatin was sonicated in Bioruptor (Diagenode) at high power for 20 min (30 sec on/off, 20 cycles). 25-50 μg sonicated chromatin were used for each IP (500 μL). 50 μl (10%) from each diluted extract for IP were taken as the inputs. For ChIP using anti-HA beads, HA IPs from WT E14 cells were used as control IPs. 20 μL of anti-HA magnetic beads (Pierce) were added to lysates from KI cells (as IPs) and E14 cells (as control IPs) for 4°C overnight incubation. For ChIP using endogenous anti-CMTR1 antibody, 2μg of anti-CMTR1 antibody (Sigma HPA029979) were added to lysates from *Cmtr1* KI cells and E14 cells. For RNAPII ChIP, 2μg of anti-Rpb1 (CST 14958) antibody were added to lysates from E14 (*Cmtr1^WT^*), *Cmtr1^KD^* or *Cmtr1^KD^* + HA-CMTR1 cells. The endogenous antibodies were added for 4 °C overnight reaction before the incubation with protein G Dynabeads (Life Technologies) for additional 2 hrs. The beads were washed and eluted as described before the proteinase K treatment and reverse crosslinking at 42°C overnight. The eluted DNA was purified by the QIAquick PCR purification kit (QIAGEN) (27).

#### RNA extraction and cDNA synthesis

RNA was extracted using TRIzol Reagent (Thermo Fisher) and concentration determined using a NanoDrop 2000 spectrophotometer. Genomic DNA was removed using the TURBO DNA-free Kit (Thermo Fisher). 400-500 ng RNA was used in 20μl cDNA synthesis reactions using iScript cDNA Synthesis Kit (Bio-Rad). Three biological replicates were used for the PCR analysis or sequencing.

#### Nuclear run-on assay

3 x10^6^ cells were seeded on a 10 cm plate one day before the experiment. Nuclear run-on was performed as previously described (27). 1 μg recombinant CMTR1 protein was added to nuclei from a 10 cm plate and incubated for 20 min at 30 °C, with 1 mM ATP, 1 mM CTP, 1 mM GTP, 0.5 mM BrUTP, 0.5 mM UTP and 1 U/mL RNAsin. After nuclear run-on and RNA extraction, Br-U IP was performed with 8-12 μg RNA. Following IP, nascent RNA was purified and subjected to cDNA synthesis and qPCR analysis.

#### qPCR

5 μL qPCR reaction contained 1 μL DNA template (undiluted IP or 10× diluted input for ChIP-qPCR and nuclear run-on qPCR or 2× diluted cDNA for RT-qPCR), 2.5 μL SsoFast EvaGreen supermix (Bio-Rad), 0.5 μL primer pairs (see list below). Each qPCR reaction was performed in triplicate on CFX384 Touch Real-Time PCR Detection System (Bio-Rad). 3 min denaturation at 95°C, followed by 40 cycles of 10 sec at 95°C, 20 sec at 58°C and 10 sec at 72°C and final incubation at 72°C for 5 min. For ChIP-qPCR, ratio of Ct value of IP to 1% input was calculated to determine ChIP efficiency. For RT-qPCR and nuclear run-on qPCR, the Ct values of target genes were normalised to corresponding Ct value of Actin and or Gapdh in each treatment condition to obtain relative transcript levels.

#### Library preparation and sequencing

For ChIP-seq, libraries were prepared using NEBNext® Ultra™ II DNA Library Prep Kit and NEBNext® Multiplex Oligos (Index Primers Set 1) for Illumina. For RNA-seq, ribosomal RNA depletion was performed on 1 μg RNA using the NEBNext® rRNA Depletion Kit v2 (Human/Mouse/Rat). RNA-seq library preparation was carried out using NEBNext® Ultra™ II Directional RNA Library Prep Kit and indexed with NEBNext Multiplex Oligos for Illumina (96 Unique Dual Index Primer Pairs). Sequencing was performed with a read length of 2×75 bp on Illumina NextSeq 500 at the Tayside Centre for Genomic Analysis.

#### Bioinformatics analysis

The quality of sequencing reads was checked using FastQC and adaptors were trimmed using Cutadapt. Trimmed reads were aligned to mouse genome assembly GRCm38/mm10 using STAR (28). Library complexity of mapped reads was assessed using preseq (29), before removal of PCR duplicates using Picard. Correlation of reads between the replicates was analysed using deepTools (30). For ChIP-seq, only uniquely mapped reads were used for data analysis. Multi-mapped reads were removed using SAMtools (31). HA IPs from WT cells were used as controls for anti-HA ChIP-seq analysis. Inputs were used as controls for endogenous RNAPII ChIP-seq analysis. For genome-wide data visualisation, metagene profiles and heatmaps were generated by ngs.plot (32). For data visualisation of individual genes, bigwig files were generated by deepTools (30) and loaded to Integrative Genomics Viewer (33). For analysis of read density near TSS, reads aligned to TSS ± 500 bp were counted using featureCounts (34). Read counts were normalized to library size and differential binding analysed using QLFTest in EdgeR to identify target genes. For gene ontology analysis, webtool Enrichr was used (35). Protein interaction networks were constructed by STRING (36). For RNA-seq, transcripts were quantified using featureCounts. Only transcripts expressed above 1 read per million in more than 3 replicates were used in the following analysis. Read counts were normalized to the library size and differential expression was analysed using exactTest in EdgeR.

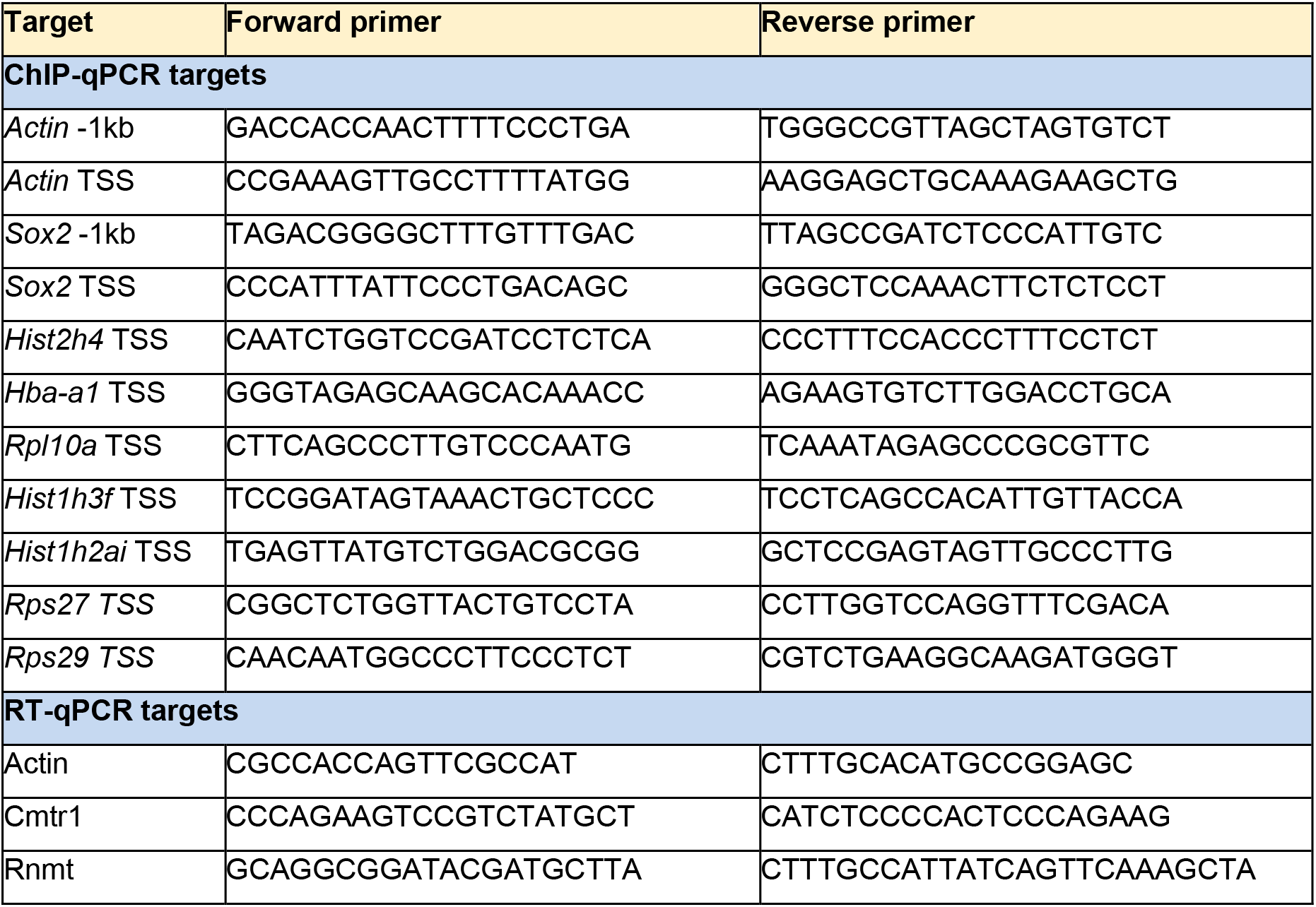
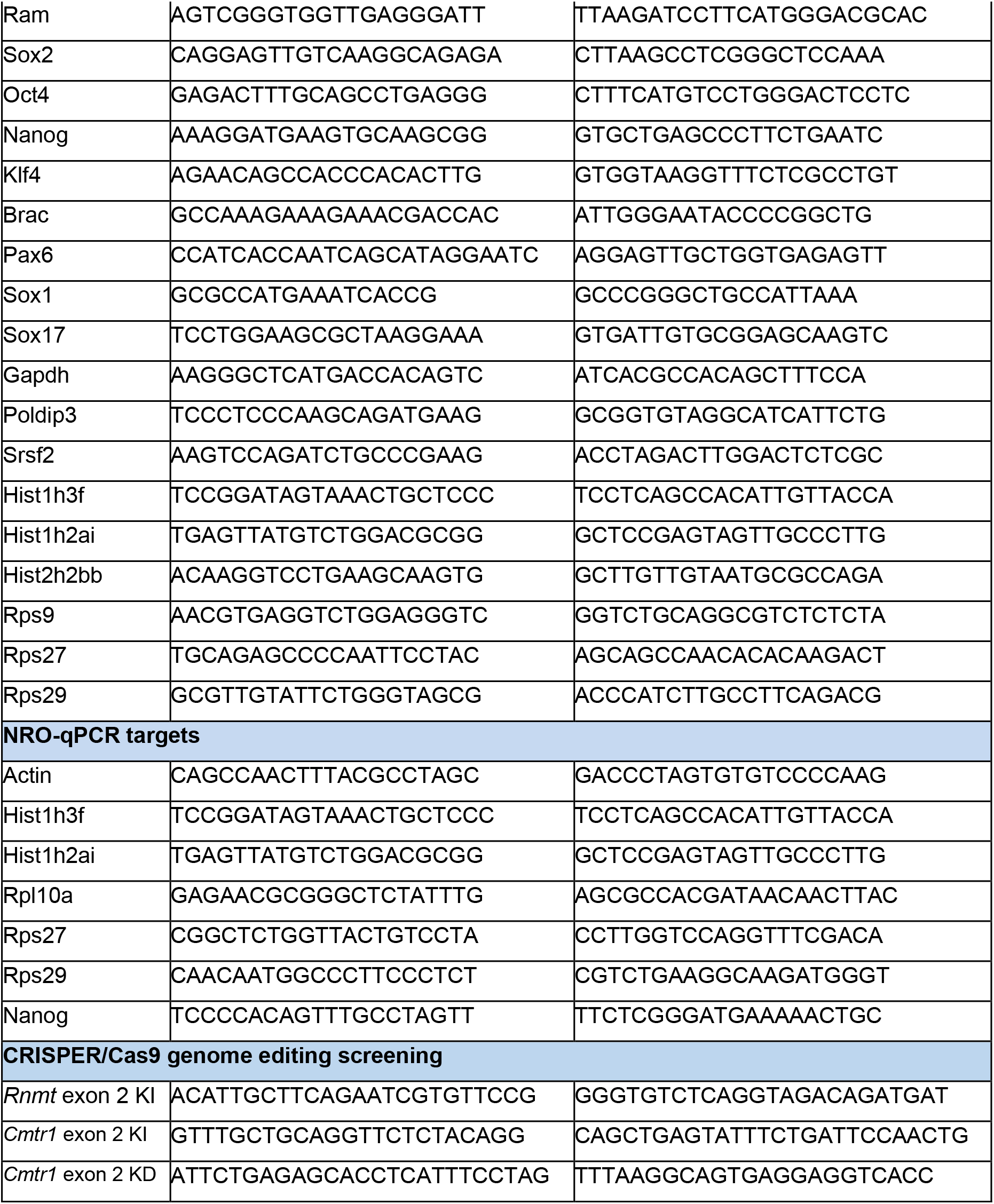

## RESULTS

### CMTR1 is recruited to the transcription start site (TSS)

We aimed to define and compare the target gene specificity of the cap methyltransferases RNMT and CMTR1 in murine embryonic stem cells (ESCs). To facilitate a comparison of the RNMT and CMTR1 genomic binding sites, CRISPR-Cas9 HA-GFP knock-in (KI) ESC lines were generated in which the endogenous *Rnmt* and *Cmtr1* genes were expressed as N-terminal HA-GFP fusion proteins (Figure S1a, b). Compared to the untagged endogenous proteins in control ESCs, HA-GFP-RNMT and HA-GFP-CMTR1 were expressed at a reduced level (Figure S1b-d). The localisation of HA-GFP-RNMT and HA-GFP-CMTR1 was assessed by fluorescence microscopy and immunofluorescence analysis to be equivalent to the endogenous proteins (Figure S2). As published previously, endogenous RNMT in ESC was nuclear and HA-GFP-RNMT was localised equivalently (26). Endogenous CMTR1 in ESC and HA-GFP-CMTR1 were also nuclear. Interaction of HA-GFP-RNMT and HA-GFP-CMTR1 with the known binding partners, RAM and DHX15, respectively, was maintained in the HA-GFP KI cells lines indicating that the tag was not interfering with complex formation (Figure S3). Although, HA-GFP-RNMT and HA-GFP-CMTR1 were expressed at reduced levels compared to the endogenous protein, in part because they were expressed from a single allele, they are not inherently unstable; neither the endogenous proteins or HA-GFP fusions were stabilised by treatment with the proteosome inhibitor, MG132 (Figure S4). Furthermore, the expression level of key pluripotency-associated transcription factors Sox2 and Klf4 were comparable in control and engineered ESCs (Figure S1c).

To compare the genomic recruitment pattern of RNMT and CMTR1 as equivalently as possible, Chromatin Immunoprecipitation-DNA sequencing (ChIP-seq) assays were performed using anti-HA antibodies to immunoprecipitated (IP) HA-GFP-RNMT and HA-GFP-CMTR1 in ESC lines, with parental ESCs used as a negative control (Figure S1d). Following DNA sequencing, CMTR1 was found to be enriched at the 5’ end of genes, peaking proximal to the TSS (transcription start site) (Figure1a, S5, S6, Table S1). More than 700 genes had significant CMTR1 binding within 500 bp of the TSS (designated “CMTR1-enriched genes”, Figure 1b, Table S1). In contrast, RNMT was not significantly enriched at genomic loci; only 20 genes, including coding and non-coding, were found to have RNMT enrichment (Figure 1b, Table S1). Most of the RNMT-enriched genes (13 out of 20) colocalised with CMTR1 near the TSS, but lower numbers of reads were produced from the RNMT ChIP, consistent with reduced genomic binding compared to CMTR1 (Figure 1c). RNMT was immunoprecipitated equivalently to CMTR1 via the HA tag (Figure S1d).

**Figure 1.**
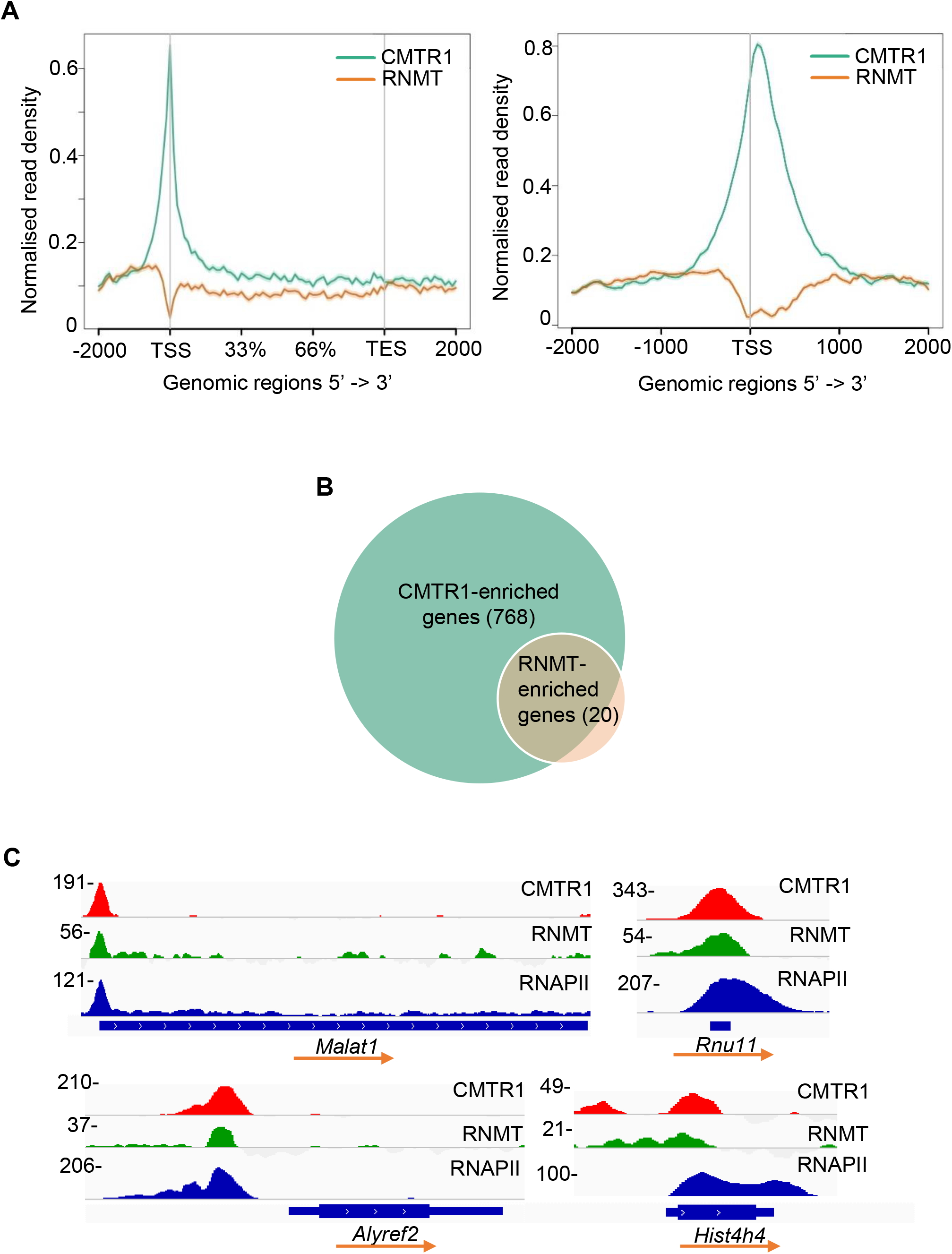
CMTR1 is recruited to transcription start sites (TSS) (A) Metaplots of normalised genome-wide CMTR1 and RNMT ChIP-seq signals ±2kb around gene body (left) and TSS (right). Read densities (reads per kilobase per million, RPKM) were normalised to library size and control IPs to obtain the fold changes in ChIP read densities relative to controls. (B) Venn diagram of number of CMTR1 and RNMT-enriched genes. Mapped reads ± 500 bp around TSS were normalised to library size to obtain RPKM. Enriched genes were defined as those with significant binding above control (fold change ≥1.5, P-value ≤ 0.05, FDR adjusted P-value ≤ 0.05). (C) Integrative Genomics Viewer (IGV) depictions of normalised CMTR1, RNMT and RNA polymerase II (RNAPII) read densities on sample genes. ChIP read densities were normalised to library size and control. Transcription direction indicated.

ChIP PCR performed with an anti-HA antibody was used to verify that HA-GFP-CMTR1 bound significantly to the TSS whereas HA-GFP-RNMT did not (Fig S7a). ChIP-PCR performed using a polyclonal anti-CMTR1 antibody was used to confirm that tagged and endogenous CMTR1 enriched equivalently at the TSS of individual genes, despite the HA-GFP-CMTR1 being expressed at a reduced level compared to the endogenous protein (Figure S7b). Since the majority of RNAPII-transcribed RNA receives a 7-methylguanosine cap during transcription, RNMT may be recruited predominantly via the guanosine cap intermediate and/or its interaction with RNA (27,37). The limited binding of RNMT to the TSS in ESC is consistent with previous work performed in HeLa cells in which RNMT had to be overexpressed for RNMT binding to the TSS to be detected (38) (see discussion).

### CMTR1 colocalises and correlates with RNAPII binding near the TSS

We analysed the identity of genes with CMTR1 enrichment using Gene Ontology (GO) analysis. The 768 CMTR1-enriched genes predominantly encoded RNA/DNA-binding proteins and were associated with biological processes which included transcription, RNA metabolism and translation (Figure 2a). The 20% most CMTR1-enriched genes were histones, ribosomal proteins and RNA processing and transport factors, indicating potential common features of these genes or their binding proteins which promote CMTR1 recruitment (Figure 2b, c).

**Figure 2.**
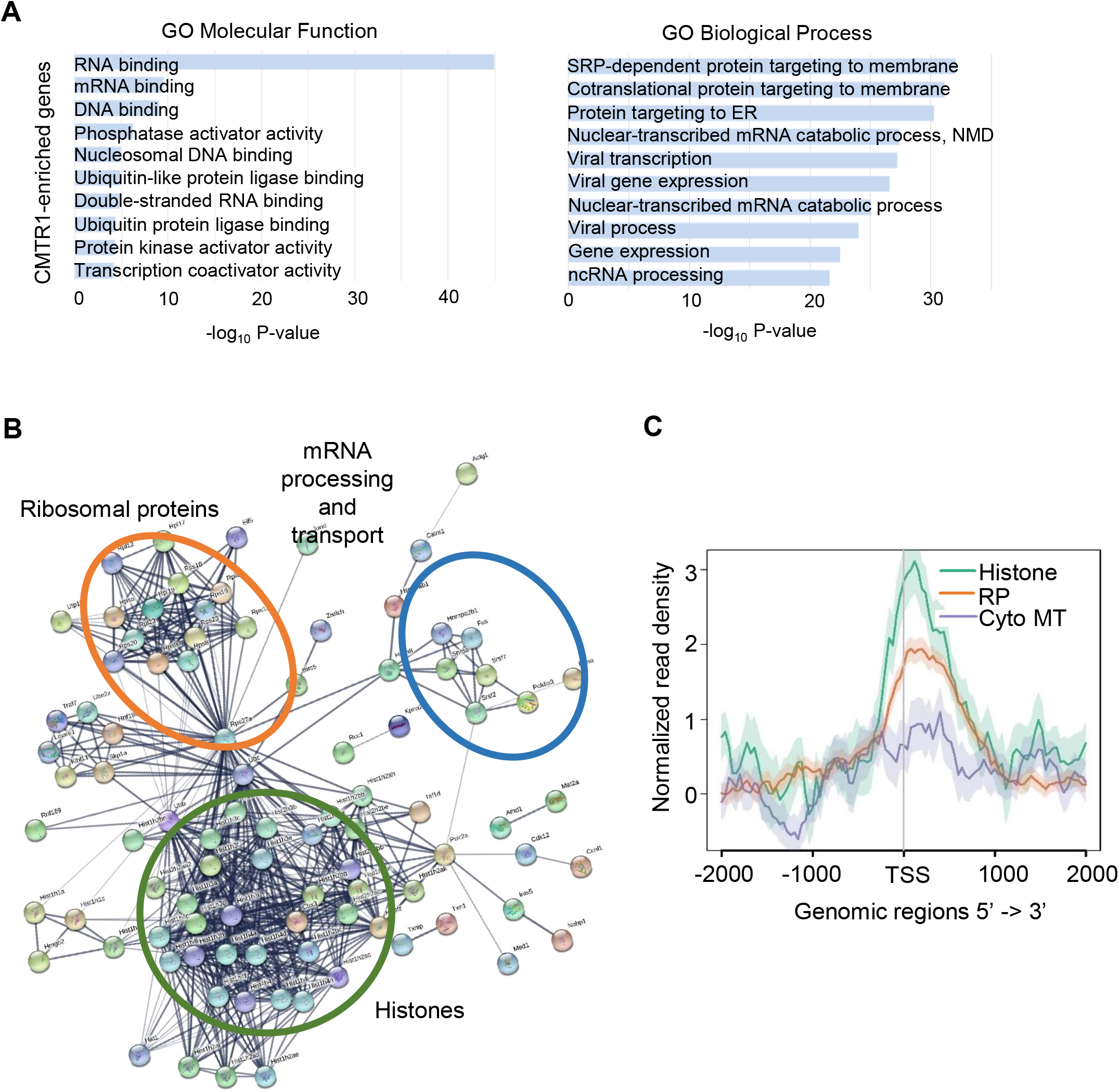
Histone and ribosomal protein genes have highest level of CMTR1 binding. (A) Gene ontology (GO) analysis of CMTR1-enriched genes (768 genes). Top 10 GO Molecular Function and GO Biological Process terms reported by significance. (B) Protein interaction network of the top 20% protein-encoding CMTR1-enriched genes (154 genes). Lines indicate established interactions from experiments and databases from STRING functional protein association networks. Disconnected nodes were hidden. (C) Metaplots of normalised CMTR1 ChIP-seq signals ±2kb around TSS in histone (68 genes), ribosomal protein (RP, 100 genes) and cytoplasmic microtubule (Cyto MT, 70 genes).

Previous studies demonstrated that CMTR1 binds directly to RNAPII CTD phosphorylated on Serine-5 (RNAPII Ser5P) (23), and therefore we investigated the correlation between RNAPII and CMTR1 chromatin binding (Figure 3, S8). RNAPII recruitment was determined by RNAPII ChIP-seq (Table S1). RNAPII Ser5P recruitment was determined using a previous dataset (39). Genome-wide, colocalisation of CMTR1 with total RNAPII and RNAPII Ser5P was observed, adjacent to the TSS, in coding and non-coding genes (Figure 3a, b). Moreover, the TSS read density in the CMTR1 ChIP samples correlated with total RNAPII and RNAPII Ser5P in expressed genes (Figure 3c, S8), consistent with CMTR1 recruitment to RNAPII Ser5P during the early stages of transcription. Notably, CMTR1-enriched genes, including histones and ribosomal proteins, displayed high promoter-proximal levels of total RNAPII and RNAPII Ser5P (Figure 3c, 3d, S8).

**Figure 3.**
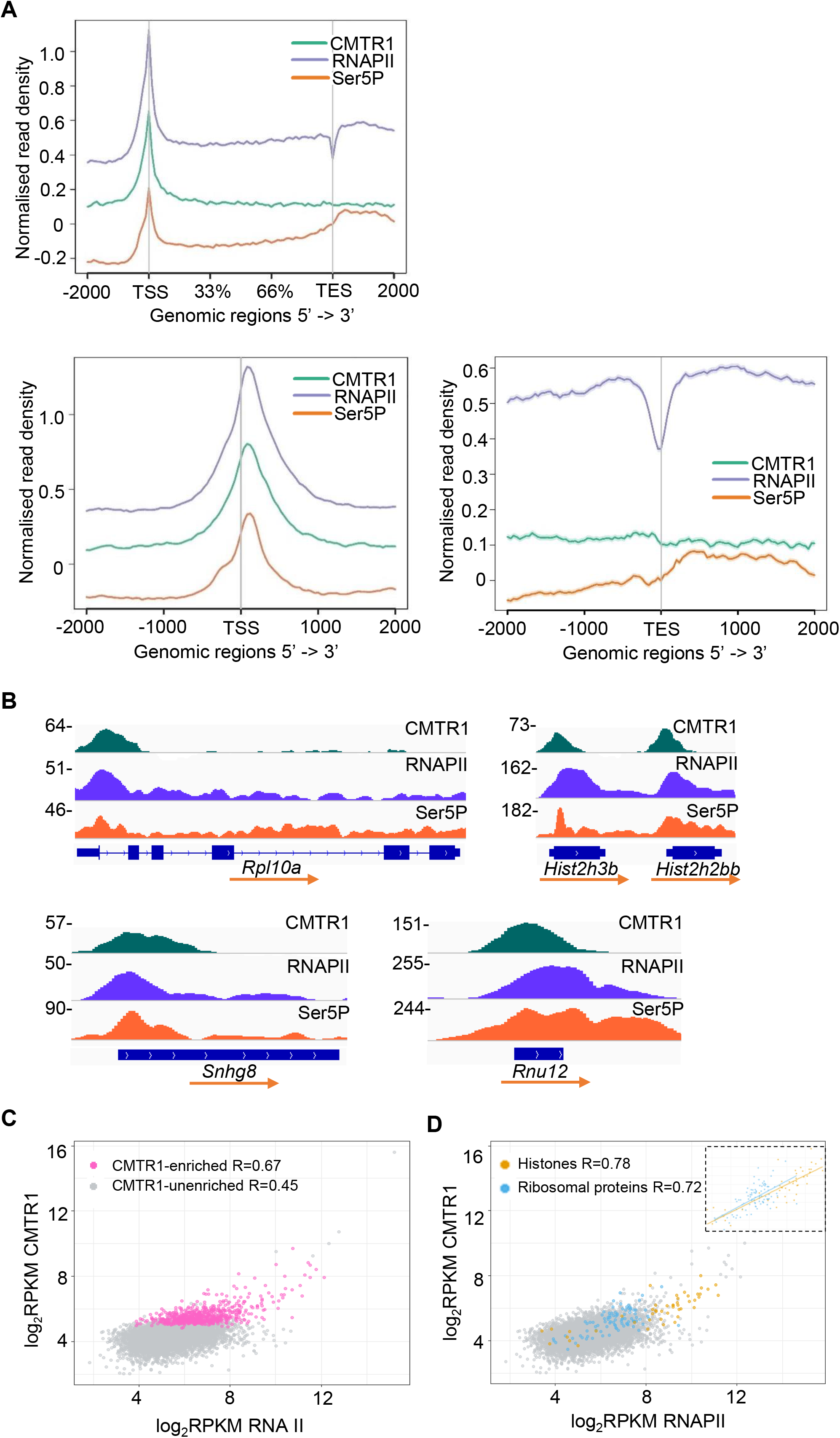
CMTR1 TSS binding correlates with RNAPII binding. (A) Metaplots of normalised genome-wide CMTR1, total RNAPII and RNAPII Ser5P CTD ChIP-seq signals ±2kb around gene body, TSS and TES. (B) IGV depictions of normalised CMTR1, RNAPII and Ser5P read densities on sample genes. (C, D) Dot plots indicating CMTR1 and RNAPII read densities (log_2_RPKM) ±500b around TSS for expressed genes. (C) CMTR1-enriched genes highlighted (pink,768 genes). (D) Histones (orange, 68 genes) and ribosomal protein (blue, 100 genes) highlighted.

In addition to RNAPII, the TSS may hold other factors which aid recruitment of CMTR1. We inspected the interactome of CMTR1 for such auxiliary factors but did not find a protein which correlated with CMTR1 binding better than RNAPII (40). The helicase DHX15 interacts with CMTR1 and influences gene expression (23,24), although we found this interaction to be minimal in ESC (Figure S3b). DHX15-CMTR1 complexes are excluded from RNAPII interactions and DHX15 does not influence the interaction of CMTR1 with RNAPII (23). The hypomethylated guanosine cap may contribute to TSS recruitment. A peak of RNAPII CTD Ser5P was also observed downstream of the transcription end site (TES), which did not correlate with CMTR1 binding (Figure 3a, S9).

### CMTR1 influences RNAPII TSS levels

Since m7G cap formation is associated with a transcription checkpoint, we investigated the impact of CMTR1 on RNAPII recruitment and retention during the early stages of transcription. RNAPII ChIP-seq was performed following knockdown of endogenous CMTR1 using siRNA. Metagene plots revealed a global reduction in RNAPII binding at the TSS and gene body (Figure 4a), indicating that CMTR1 was influencing RNAPII recruitment or retention. In single gene analysis, out of genes with detectable TSS RNAPII binding, 7299 genes had a reduction in promoter-proximal RNAPII levels following CMTR1 siRNA transfection, whereas 3385 genes had increased RNAPII (Figure 4b, 4c, Table S1).

**Figure 4.**
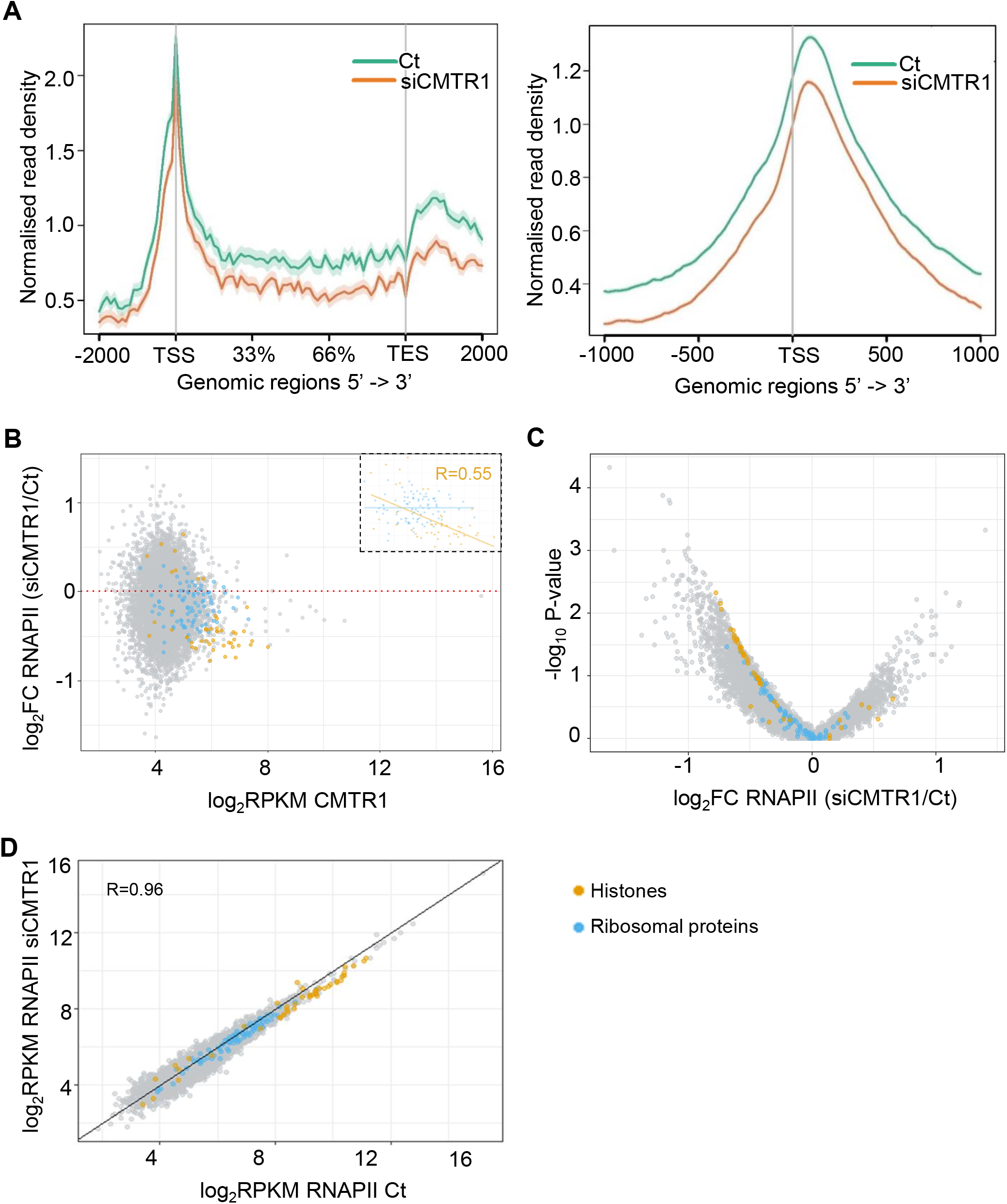
A reduction in CMTR1 expression results in loss of RNAPII TSS binding. ESCs were transfected with CMTR1 siRNA 1 (siCMTR1) or control (Ct) for 36h. (A) Metaplots of normalised genome-wide RNAPII ChIP-seq signals in gene body ±2kb (left) and TSS ±1kb (right). (B) Dot plot of CMTR1 read densities (log_2_RPKM) in control ESC and log_2_ fold change in RNAPII ChIP signal (TSS ±500b) following transfection with CMTR1 siRNA 1. (C) Volcano plot indicating the relationship between log_2_FC and -log_10_ FDR adjusted P-value in RNAPII ChIP (TSS ±500b) following transfection with CMTR1 siRNA 1. (D) Dot plot indicating the relationship between RNAPII binding in cells transfected with CMTR1 siRNA and control. Histones (orange) and ribosomal protein (blue) genes are labelled.

Although a reduction in CMTR1 expression resulted in reduced average RNAPII binding, for single genes the decrease in RNAPII binding did not correlate with TSS CMTR1 levels (Figure 4b). However, for the two groups of genes with the highest CMTR1 binding, the histones and ribosomal protein genes, following CMTR1 repression RNAPII TSS binding was predominantly repressed (Figure 4b, c). In particular for histone genes, the decline in TSS RNAPII levels correlated with CMTR1 binding indicating a potential role for CMTR1 in regulating their transcription directly (Figure 4b). Inspecting RNAPII binding to individual genes in cells transfected with CMTR1 siRNA or control confirmed these findings (Figure 4d). The histone genes had some of the highest levels of RNAPII binding and this was reduced for almost all when CMTR1 was repressed. RNAPII binding to ribosomal protein genes was also largely reduced following CMTR1 repression. Overall, there was a good correlation of RNAPII binding to other genes in cells transfected with CMTR1 or control siRNA, indicating that CMTR1 is not influencing RNAPII recruitment to the majority of genes.

### CMTR1 has gene specific impacts on transcript levels

Since CMTR1 repression resulted in changes in TSS RNAPII levels, we investigated the impact on the transcriptome by performing RNA sequencing analysis on ESCs in which CMTR1 had been repressed by transfection of CMTR1 siRNAs (Figure S10, Table S1). In control ESCs, there were correlations between TSS CMTR1 binding and transcript levels for the histone and ribosomal protein genes (Figure S11). For the CMTR1-enriched genes, ~60% were down-regulated following CMTR1 knockdown (Figure 5a). The genes with the highest CMTR1 binding, encoding histones and ribosomal proteins, were significantly repressed following CMTR1 repression, consistent with reduced RNAPII binding (Figure 5b, 4b). 87/100 ribosomal protein genes and 54/68 histone genes were repressed following CMTR1 knock-down (Table S1). The correlation coefficients between fold change in transcript level and CMTR1 binding in histone and ribosomal proteins were most significant for these gene sets (Figure 5b). In addition, the fold change in transcript level following CMTR1 repression also correlated with RNAPII TSS binding, for the histone and ribosomal protein genes (Figure S12). For controls, we analysed some highly expressed but CMTR1-unenriched genes, including pluripotency-associated and splicing-associated genes. The fold changes of these transcripts did not correlate with CMTR1 or RNAPII binding, suggesting that the influence of CMTR1 on these genes is likely indirect (Figure 5c, 5d, S11, S12).

**Figure 5.**
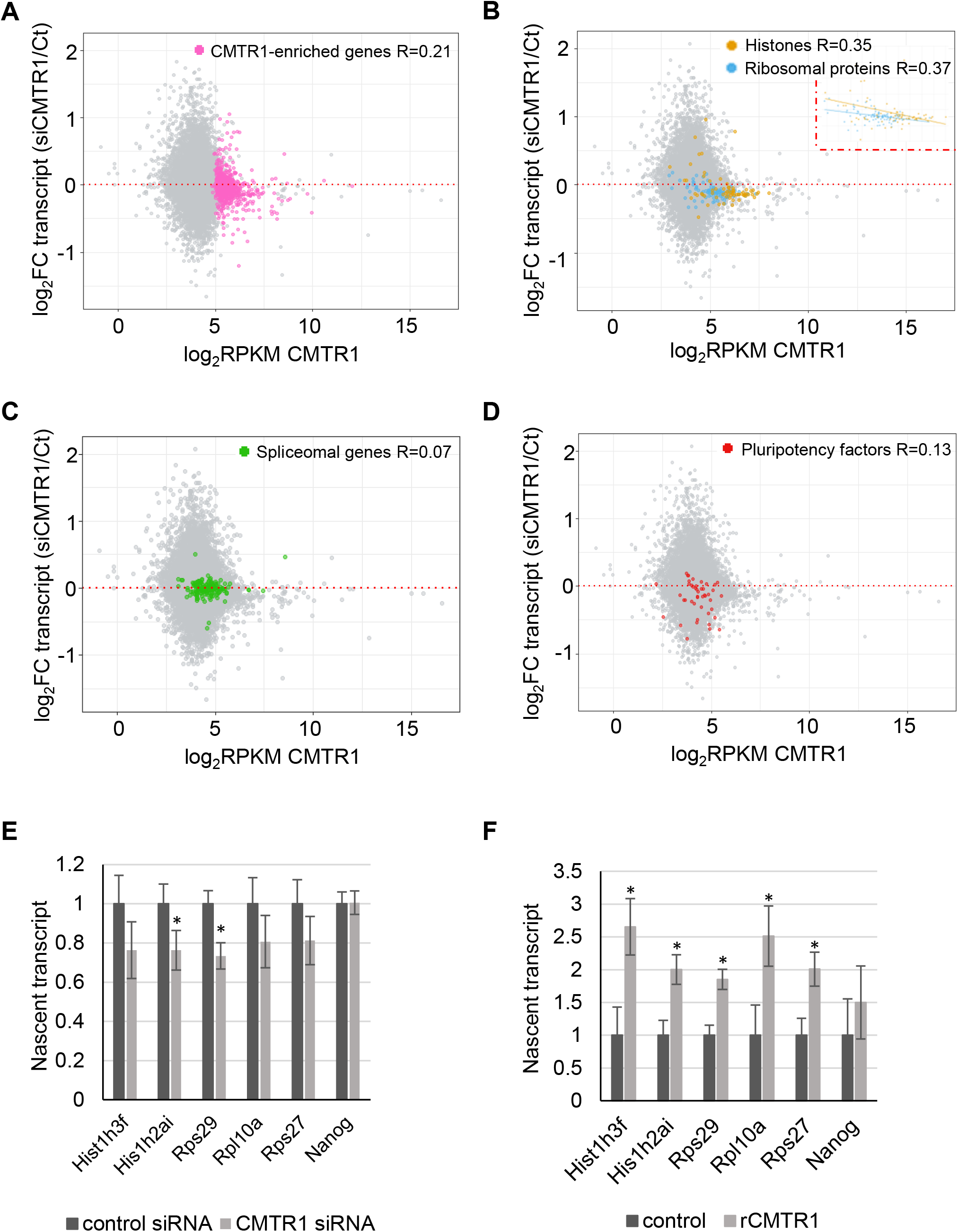
CMTR1 promotes expression of ribosomal protein and histone genes. (A-D) Dot plots of CMTR1 binding (TSS ±500b) in control cells and changes in RNA levels following CMTR1 siRNA transfection. Highlighted: (A) CMTR1-enriched genes (768 genes), (B) histones genes (68 genes) and ribosomal protein genes (100 genes), (C) spliceosomal genes (154 genes) and (D) pluripotency genes (45 genes). Pearson correlation coefficients (R) indicated. (E, F) RT-qPCR analysis of nascent RNA following nuclear run-on assay (N=3). (E) Nuclei were prepared from ESC transfected with CMTR1 siRNA or control. (F) recombinant (r) CMTR1 protein was added to ESC nuclei. For each replicate, RNA level was normalised to the corresponding nascent Actin RNA level. For charts, average and standard deviation of three experiments presented. “*” indicates a P-value <0.05 for the Students T test, performed relative to control.

Since we observed that the histones and ribosomal protein genes have the highest levels of CMTR1 recruitment (Table S1, Figure 2b, 3d), and RNAPII and transcript levels were significantly repressed on CMTR1 repression (Figure 4b, 5b), we investigated whether their transcription can be directly influenced by CMTR1. Nascent transcription of histone genes and ribosomal protein genes in isolated ESC nuclei was analysed in nuclear run-on assays (Figure 5e, f). Nascent transcription of histone genes and ribosomal protein genes was reduced in ESCs in which CMTR1 was repressed (Figure 5E). Conversely, addition of recombinant CMTR1 to ESC nuclei stimulated nascent transcription of histone and ribosomal protein genes (Figure 5f). As a control, transcription of the pluripotency factor Nanog was investigated, which was not significantly regulated by CMTR1 repression or addition (Figure 5E and F). These results confirmed that CMTR1 can have a direct impact on nascent transcription of histone and ribosomal protein genes.

### CMTR1 is upregulated during neural differentiation of ESC

To determine the biological impact of CMTR1, we investigated its role in ESCs in the pluripotent state and during differentiation. To induce differentiation, the growth factor LIF (Leukemia Inhibitory Factor) was withdrawn from the medium resulting in morphological changes associated with differentiation, suppression of pluripotency factors (Oct4, Nanog, Sox2 and Klf4), and an increase in markers of differentiation (Brachyury, Sox17 and Sox1) (Figure 6A). During LIF withdrawal-induced differentiation, CMTR1 protein was upregulated independently of changes in CMTR1 RNA level, indicating a post-transcriptional mechanism (Figure 6B). CMTR1 is likely to be upregulated either by increases in translation rate and/or decreases in protein turnover. ESCs were also differentiated towards a neural fate by culture in N2B27 medium, with associated repression of pluripotency factors and induction of the neural marker, Pax6 (Figure 6c). CMTR1 protein was also upregulated by a post-transcriptional mechanism during neural differentiation (CMTR1 RNA level was maintained) (Figure 6d).

**Figure 6.**
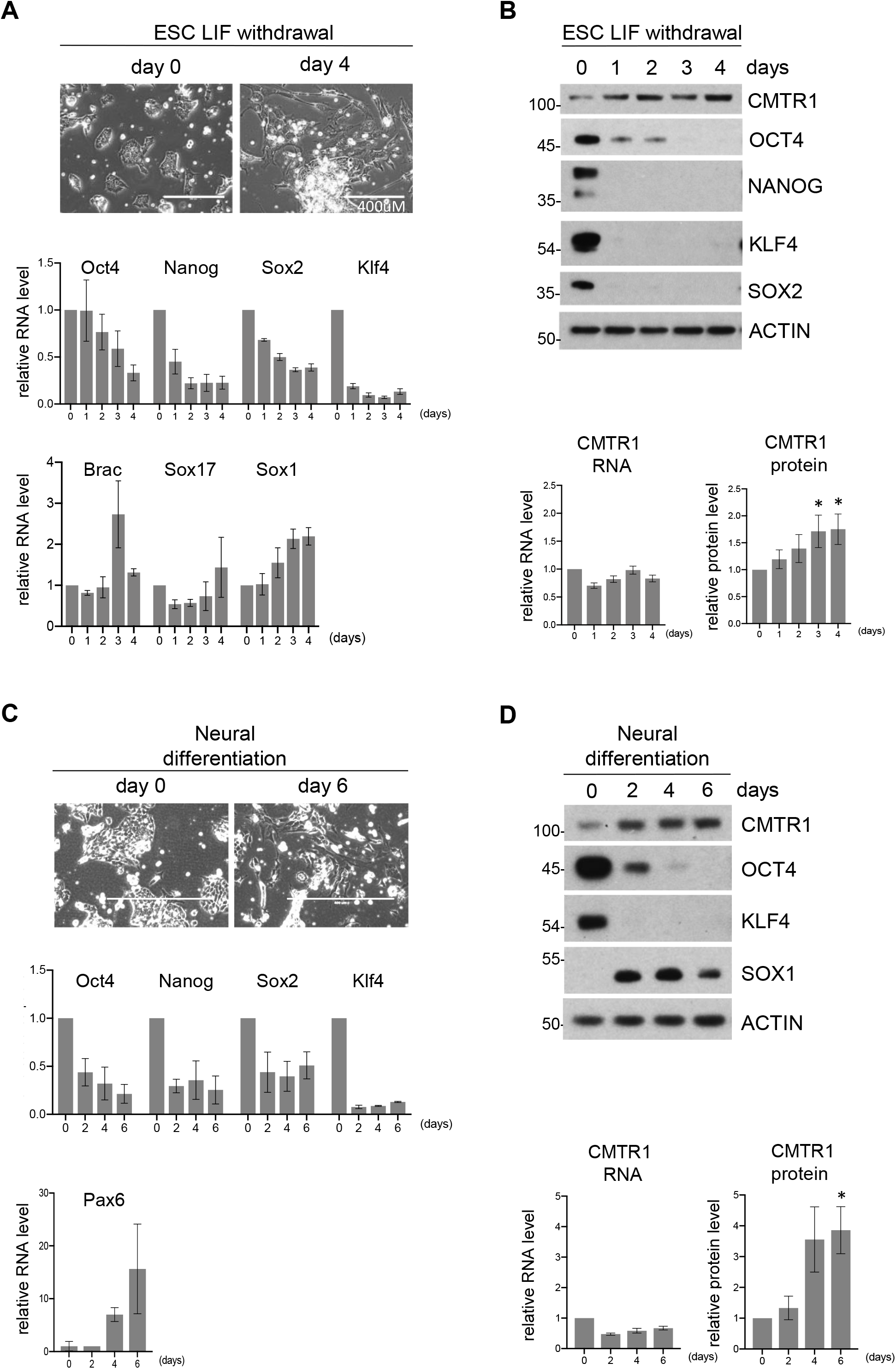
CMTR1 protein expression increases during ESC differentiation. Differentiation was induced in ESC by LIF withdrawal. (A) Phase contrast images (top) and RT-PCR analysis of pluripotency-associated (Oct4, Nanog, Sox2, Klf4) and differentiation-associated (Brac, Sox17, Sox1) genes presented (bottom). (B) Representative western blot analysis of CMTR1 and pluripotency associated proteins (top) and quantification of CMTR1 RNA and protein levels (bottom). Neural differentiation was induced in ESC. (C) Phase contrast images (top) and RT-PCR analysis of pluripotency-associated (Oct4, Nanog, Sox2, Klf4) and neural (Pax6) genes presented (bottom). (D) Representative western blot analysis of CMTR1 and pluripotency associated proteins (top) and quantification of CMTR1 RNA and protein levels (bottom). For charts, average and standard deviation of 3+ experiments presented. For CMTR1 protein chart, “*” indicates Student’s T test P-value <0.05. RNA levels were normalised to *Actin* and *Gapdh* and relative to day 0. Proteins levels were quantified relative to day 0 and normalised to Actin by densitometry using ImageJ.

To investigate the role of CMTR1 in differentiation, we used CRISPR technology in an attempt to delete the *Cmtr1* gene. Despite taking several approaches, we were unable to create a *Cmtr1* knock-out cell. However, we were able to achieve mutation of the *Cmtr1* gene in which 6bp were deleted and 2 bp were mutated; this created an in-frame deletion of 2 amino acids, Ser 30 and Asp31, and mutation of Asp32 to Cys32 and Glu33 to Val33. We named this mutant *Cmtr1^KD^* because the expression of CMTR1 was significantly reduced (Figure 7a, S13). The *Cmtr1^KD^* mutation is not predicted to have a major impact on CMTR1 protein function because it is in a disordered region which is not part of the catalytic domain or a site of known protein: protein interaction (14,15,23). CAP-MAP mass spectrometry analysis of mRNA in the *Cmtr1^KD^* cell line indicated a reduction in the mature caps, ^m7^GpppG_m_, ^m7^GpppA_m_ and ^m7^Gppp^m6^A_m_ and an increase in the immature caps lacking N1 2’-O methylation, ^m7^GpppG, ^m7^GpppA and ^m7^Gppp^m6^A compared to control ESCs (Figure 7b)(4). This reduction in N1 2’-O-Me is consistent with a reduction in CMTR1 expression and indicates the absence of a completely redundant methyltransferase for this modification.

**Figure 7.**
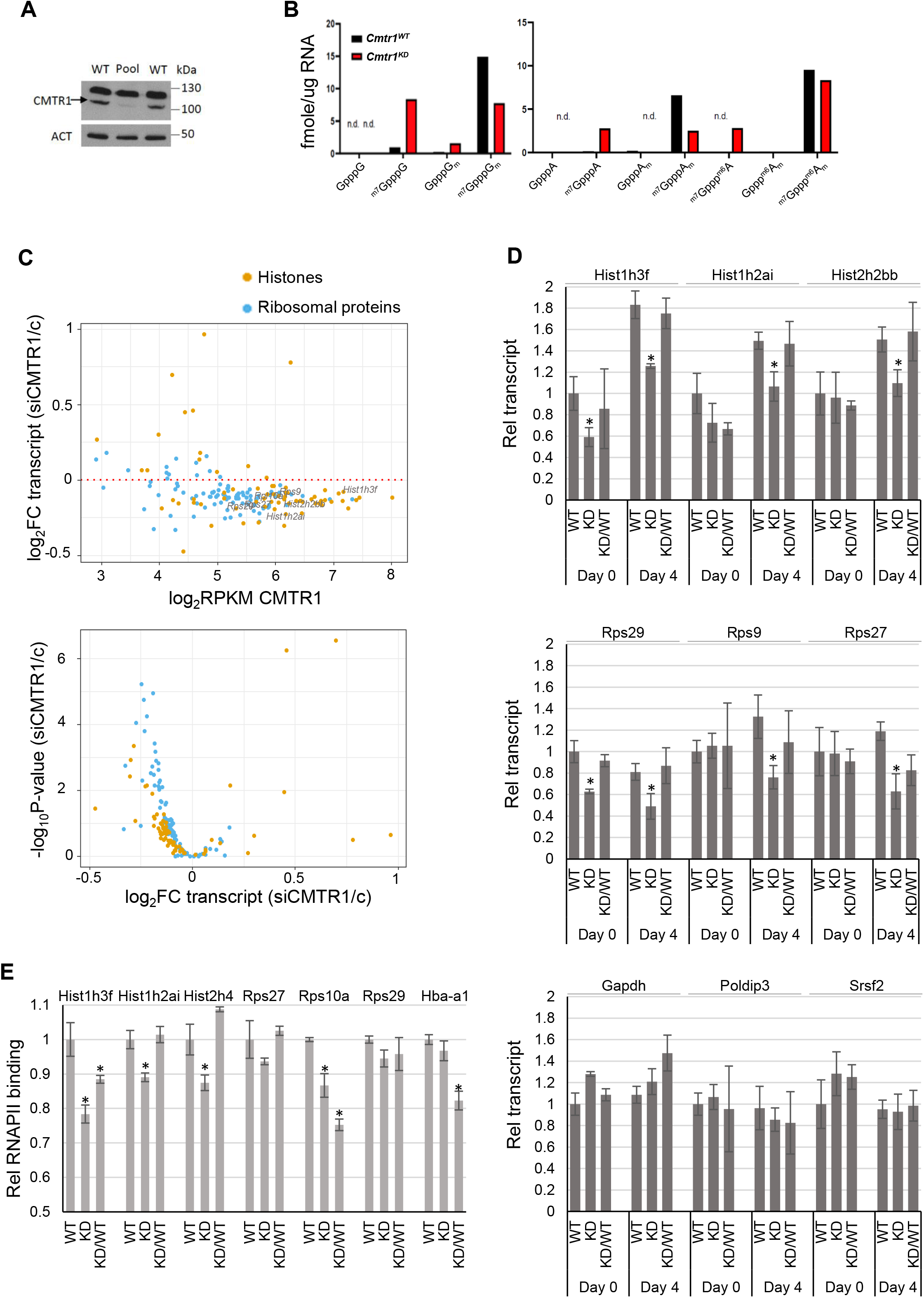
CMTR1 is required for histone and RP gene expression. (A) Western blot analysis of CMTR1 protein expression in *Cmtr1^WT^* and *Cmtr1^KD^* ESCs. Band verified by CMTR1 IP and KO. (B) CAP-MAP RNA cap structure analysis in *Cmtr1^WT^* and *Cmtr1^KD^* ESCs. “N.d.”, indicates not detected. (C) Regulation of histone and ribosomal protein genes following CMTR1 siRNA transfections (siCMTR1) in ESCs was determined by RNA seq analysis. Dot plots indicates transcript level in control cells and Log_2_FC following CMTR1 siRNA transfection. Volcano plots demonstrates Log_2_FC expression and EdgeR exactTest -log_10_ FDR adjusted P-value. (D) RT-PCR analysis of genes indicated relative to Actin RNA in *Cmtr1^WT^, Cmtr1^KD^* and *Cmtr1^KD^/*HA-CMTR1 ESCs, in cells cultured in LIF (day 0) and neuro differentiated cells (day 4). RNA levels were normalised to *Actin* and relative to day 0. (E) RNAPII ChIP-PCR in *Cmtr1^WT^,Cmtr1^KD^* and *Cmtr1^KD^/HA-CMTR1* ESCs. Unexpressed gene *Hba-a1* was used as a negative control. D, E) N=3. Average and standard deviation presented. “*” Student’s T test P-value <0.05, relative to control.

### CMTR1 is required for histone and ribosomal protein gene expression during differentiation

Initially we investigated whether there was concordance between observations made in CMTR1 siRNA treated cells and *Cmtr1^KD^* ESCs. Following CMTR1 siRNA transfection there is a rapid reduction in CMTR1, whereas in *Cmtr1^KD^* ESCs there is long-term, minimal CMTR1 expression. As established previously, histone and RP RNA expression was reduced on CMTR1 siRNA transfection (Figure 7c, S14, Table S1). In the *Cmtr1^KD^* ESC, there was also a reduction in the histones and RPs, Hist1h3f, Hist1h2ai and Rps29, compared to *Cmtr1^WT^* ESCs, when cultured in LIF (Day 0) (Figure 7d, S15). Other histones and RPs were not repressed in *Cmtr1^KD^* ESC when cultured in LIF (Hist2h2bb, Rps9, Rps27). On neural differentiation of *Cmtr1^KD^* ESC, all histones and RP genes investigated were now repressed compared to the controls, suggesting that there is an increased requirement for CMTR1 as cells differentiate and CMTR1 is upregulated (see discussion). Expression of HA-CMTR1 in the *Cmtr1^KD^* ESC was sufficient to rescue histone and RP expression. Three control genes were not significantly regulated in the *Cmtr1^KD^* ESC; Gapdh, Srsf2, Poldip3 (Figure 7d). Confirming that there is a transcriptional component to CMTR1-regulated RNA expression, ChIP-PCR was used to confirm that RNAPII binding to histone and RP genes is also largely repressed in *Cmtr1^KD^* ESC compared to controls, and expression of HA-CMTR1 resulted in rescue of RNAPII binding for most genes investigated (Figure 7e).

To investigate the impact of CMTR1-dependent RP gene expression, net translation rates were approximated by determining the rate of ^35^S cysteine and methionine incorporation into cellular proteins. In *Cmtr1^KD^* ESC, translation was significantly reduced compared to *Cmtr1^WT^* ESC and this could be rescued by expression of HA-CMTR1 (Figure 8a). A reduction in histones may lead to DNA replication stress and damage (41,42). A marker of DNA replication stress, phospho-Chk1, was upregulated on repression of CMTR1 (Fig 8b). During differentiation, a marker of DNA replication stress and damage, gamma-H2AX, was also upregulated on repression of CMTR1 (Figure 8c).

**Figure 8.**
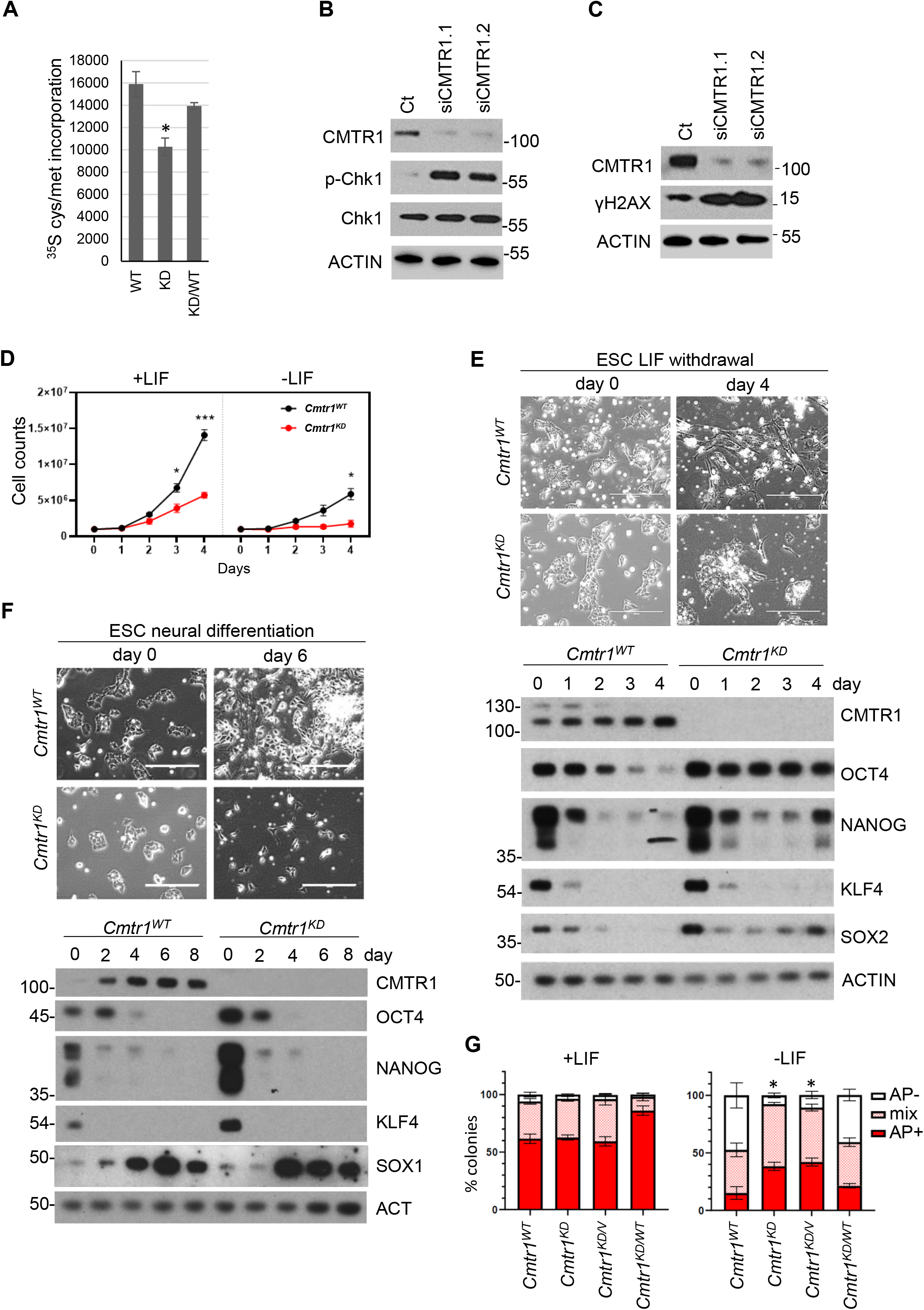
CMTR1 is required for histone and ribosomal protein gene expression during neural differentiation. A) *Cmtr1^WT^, Cmtr1^KD^* and *Cmtr1^KD^/*HA-CMTR1 ESCs were cultured in ^35^S cysteine/methionine for 30 mins and incorporation into cellular proteins was determined (N=3). Average and standard deviation is presented. Student’s t-test; “*” indicates P-value <0.05 relative to control. B, C) Western blot analysis of ESC cultured in B) LIF and C) without LIF for 2 days following CMTR1 siRNA transfection. N=3. D) *Cmtr1^WT^ and Cmtr1^KD^* ESC were cultured in and without LIF for the days indicated. Cell number was counted. N=3. Average and standard deviation presented. Student’s t-test; “*” indicates P-value <0.05 relative to control timepoint. E) *Cmtr1^WT^* and *Cmtr1^KD^* ESC were induced to differentiate E) by LIF withdrawal, F) towards a neural fate. Phase contrast images and representative western blot analysis. G) The ESC lines indicated were cultured without LIF for 6 days, then stained with alkaline phosphatase. Resultant colonies were scored as alkaline phosphatase (AP) negative (-), positive (+) or mixed. N=4. Average and standard deviation presented. Student’s t-test performed on data from AP-colonies; “*” indicates P-value <0.05 relative to control timepoint.

### CMTR1 is required for proliferation during differentiation

Since a reduction in protein synthesis and an increase in DNA replication stress is observed in *Cmtr1^KD^* ESC, we investigated the impact of CMTR1 repression on cell proliferation, a process dependent on histones for DNA replication and RPs for protein synthesis. When cultured in LIF, there was a reduction in cell proliferation in *Cmtr1^KD^* ESC and this could be rescued by expression of HA-CMTR1 (Figure 8d, S15B). When LIF was removed to initiate differentiation, *Cmtr1^KD^* ESC proliferation was further reduced, accompanied by apoptosis (Figure 8d, 8e, S15b, S15c). Cell proliferation was also severely attenuated in *Cmtr1^KD^* ESC undergoing neural differentiation (Fig 8f).

Since defects in gene expression and proliferation resulting from CMTR1 repression were exaggerated during differentiation, we investigated the regulation of pluripotency factors and markers during this process. The expression of pluripotency factors was repressed on CMTR1 siRNA transfection, however on long term inhibition of CMTR1 in *Cmtr1^KD^* ESC, pluripotency factors were increased (Figure S16a, 8e, 8f). It appears that compensatory mechanisms are occurring to maintain pluripotency factor expression on stable repression of CMTR1. However, following LIF withdrawal, *Cmtr1^KD^* ESC exhibited repression of pluripotency factors, equivalent to WT ESCs, although Oct4 was only mildly reduced (Figure 8E, 16b). 4 days into the differentiation programme, the *Cmtr1^KD^* ESCs re-expressed pluripotency factors, and expression of HA-CMTR1 rescued this phenotype (Figure 8e, S15a, s16b). For the few *Cmtr1^KD^* cells which survived induction of differentiation (Figure 8f, micrographs), the markers of pluripotency, Oct4, Nanog, Klf4, were repressed on day 4 equivalently to WT cells and the intermediate neural marker, Sox1, was upregulated equivalently to WT cells (Figure 8f). This re-expression of pluripotency factors in the differentiating *Cmtr1^KD^* ESC culture is consistent with the differentiating cells having reduced proliferation and undergoing apoptosis (Figure S15b, c). To validate these findings, we induced neural differentiation and found that *Cmtr1^KD^* ESC were also unable to proliferate during differentiation (Figure 8f).

To further investigate the role of CMTR1 in ESC proliferation and differentiation, an Alkaline phosphatase (AP) colony assay was performed. AP staining is used as a marker of pluripotency during differentiation; when LIF is withdrawn and ESCs differentiate, there is a gradual increase in cell colonies which are negative for AP staining (Figure 8g, S17) (43). In *Cmtr1*^WT^ ESCs, over a titration of LIF withdrawal, AP negative colonies increased, whereas in *Cmtr1*^KD^ ESC, AP negative colonies were minimal (Figure S17). Expression of HA-CMTR1 rescued this defect in differentiation (Figure 8g). This confirmed that there are differentiation defects in *Cmtr1^KD^* ESC.

## DISCUSSION

First transcribed nucleotide ribose O-2 methylation (N1 2’-O-Me) is present on mRNA in mammalian cells and by labelling cellular RNA as self, it protects it from degradation by innate immunity pathways (3,44). The contribution of N1 2’-O-Me to cap function in gene expression is less well defined. Here we report that a key function of the N1 2’-O-Me methyltransferase, CMTR1, in embryonic stem cells (ESCs) is to promote histone and ribosomal protein gene expression. We found that CMTR1 is recruited to the TSS, with highest binding at histone genes, ribosomal protein genes and mRNA processing and transport genes. Repression of CMTR1 using siRNA transfections or genome editing resulted in a reduction in RNAPII binding to histone and ribosomal protein genes and a reduction in their expression. Thus, we observed that CMTR1 controls RNA levels in a gene specific manner in ESCs, correlating with RNAPII binding and expression level. In neurons, CMTR1 was found to control expression of Camk2a (21), and in the innate immune response, CMTR1 was found to be required for upregulation of certain interferon stimulated genes (20). In both scenarios and presented here in ESCs, CMTR1 is controlling the expression of some of the most highly expressed genes in the cell.

To investigate the impact of CMTR1 in ESCs, lines were created with minimal CMTR1 expression (CMTR1^*KD*^). These *Cmtr1^KD^* cells had reduced proliferation when cultured in LIF, but when induced to differentiate, either by LIF removal or under conditions of neural differentiation, proliferation was severely attenuated compared to controls and cell differentiation was incomplete. As discussed, some ribosomal protein genes and histone genes were repressed in pluripotent conditions (in LIF), but they were more severely and consistently repressed during differentiation. Why ESCs exhibit enhanced dependency on CMTR1 for gene expression during differentiation is not clear, but it may be that the condensation of chromatin during differentiation requires increased CMTR1 to maintain interaction with RNAPII. Of note, the marker of DNA replication damage and stress, gamma-H2AX, was only upregulated on CMTR1 repression during differentiation, reinforcing that differentiation exaggerates the requirement for CMTR1.

Which genes mediate the proliferation defect in ESCs? Here we have established that the histone and RP genes are the most CMTR1 -dependent and their CMTR1 dependency correlates with an impact on DNA replication stress and damage, and RNA translation, respectively. Although we have not conclusively proven that the defects in histone and RP genes are responsible for the proliferative defect observed when CMTR1 is repressed (it is not possible to restore expression of such a large complement of genes), it is likely that repression of these genes and the correlating defects in protein synthesis and DNA replication make a major contribution to the proliferative defect. For example, the ~40% reduction in protein synthesis is Cmtr1^KD^ cells is unlikely to support the protein synthesis demands of cell proliferation. In addition, many other genes are CMTR1 responsive (Table S1), which may contribute to the proliferative defect in *Cmtr1^KD^*

In ESCs, the pluripotency factors were repressed following transient repression of CMTR1 using siRNA, but there was little correlation between CMTR1 binding and level of repression following CMTR1 knockdown. In addition, the Nanog gene was unresponsive to CMTR1 in the nuclear run-on assay emphasising that its dependency on the enzyme is likely to involve an indirect mechanism. In *Cmtr1^KD^* ESCs, the pluripotency factors were elevated, potentially in a compensatory mechanism. During differentiation of *Cmtr1^KD^* cells induced by LIF withdrawal, most pluripotency genes were repressed with normal kinetics during initial stages of differentiation but were re-expressed in the differentiating culture in the small population of cells which escaped the proliferation defect and apoptosis. The emergence of AP negative colonies during differentiation was also repressed in *Cmtr1^KD^* cells, indicating a defect in differentiation. During neural differentiation, the attenuation of proliferation was severe, with few cells surviving.

The impact of CMTR1 and N1 2’-O-Me on RNA levels is likely to result from direct and indirect mechanisms. For histones and ribosomal protein genes, the more CMTR1 is bound to the TSS, the more the RNA is repressed in response to CMTR1 knockdown, indicating a relatively direct mechanism connecting CMTR1 and RNA levels. Nuclear run-on assays were used to focus on the role of CMTR1 in transcription. For the ribosomal protein genes and histone genes, the impact of CMTR1 on transcript appeared direct; in isolated nuclei CMTR1 knockdown results in a reduction in their transcription and addition of recombinant CMTR1 increases their transcription. RNMT and its product the ^m7^G cap are linked to a transcriptional checkpoint involving co-transcriptional stability and protein:protein interactions (1,27,37). For CMTR1 and its product N1 2’-O-Me, it is likely that analogous influences on co-transcriptional stability and transcriptional mechanisms are governing CMTR1-dependent mRNA levels. In ESCs, the expression of other genes, including pluripotency factors, is also repressed in response to CMTR1 repression. Of note, we found that genes involved in mRNA expression, processing and transport are CMTR1 targets in ESCs, which are likely to have indirect impacts on mRNA levels.

We report that the other major cap methyltransferase, RNMT, also binds at the TSS but it was only detected above significance thresholds at 20 genes. Unlike the other capping enzymes, RNMT does not have a CTD interaction domain and a direct interaction with RNAPII has not been demonstrated (22,45–48). Since the majority of nascent RNA has a ^m7^G cap, and a redundant N-7 cap guanosine methyltransferase is not observed in RNMT depleted extracts, RNMT must be recruited to nascent guanosine caps during transcription via a different mechanism (37). A previous study demonstrated that RNMT binds to RNA in correlation with the expression of that gene, without apparent preference for motifs or RNA regions (26). RNMT is more highly expressed than the other capping enzymes and has much higher activity in *in vitro* assays; this and its affinity for RNA may waive the requirement for a CTD interaction motif on which the other capping enzymes are dependent to facilitate interaction with the guanosine cap substrate (26,27). RNMT may act as many other enzymes; a transient interaction with the guanosine cap substrate, facilitated by an affinity for the TSS and associated proteins/RNAs, is sufficient for N-7 guanosine cap methylation to occur.

The majority of mRNA has a ^m7^GpppX_m_ cap in ESCs and therefore the action of both CMTR1 and RNMT appears to be required to produce stable mRNA and viable cells (4,5). However, CMTR1 and RNMT have different target genes and quite distinct biological roles in ESCs. Whereas CMTR1 is upregulated during differentiation and is required for histone and RP gene expression and proliferation, RNMT-RAM (active RNMT complex) is repressed during differentiation and this repression is required for the process of differentiation (27). In ESCs, high levels of RNMT-RAM are required for pluripotency-associated gene expression and repression of RNMT-RAM is required for repression of these genes during differentiation (27). These differences in RNMT and CMTR1 function during differentiation may be facilitated by differences in their mechanism of recruitment to their substrate, guanosine-capped RNA, which direct differences in their major RNA targets. The differential regulation of RNMT and CMTR1 during ESC differentiation directs them to control different genes and biological pathways during this process.

## AVAILABILITY

All genomic and transcriptomic data described here are deposited in Gene Expression Omnibus under accession number GSE175631

## ACKNOWLEDGEMENT

We thank the Cowling lab and Alabert lab members for advice, assistance and reagents. We thank the Tayside Centre for Genomic Analysis and the MRC PPU Reagents and Services for CRISPR strategy design.

## TABLE AND FIGURES LEGENDS

**Table S1. Summary of ChIP-seq and RNA-seq analysis of individual genes** For CMTR1 siRNA RNA-seq, analysis from two siRNAs were shown, genes with significant changes in their transcript levels after CMTR1 siRNA knockdown were labelled (fold change ≥1.5 or ≤0.67, FDR adjusted P-value ≤ 0.05). Significantly down-regulated genes were labelled as −1, significantly up-regulated genes were labelled as 1, genes without significant changes were labelled as 0.

For CMTR1 and RNMT ChIP-seq, CMTR1 or RNMT-enriched genes were defined as those genes that had ChIP enrichment fold change ≥1.5, FDR adjusted P-value ≤ 0.05. For siCMTR1 RNAPII ChIP-seq, those genes with significant changes in their RNAPII levels were labelled (fold change ≥1.5 or ≤0.67, FDR adjusted P-value ≤ 0.05). Significantly down-regulated genes were labelled as −1, significantly up-regulated genes were labelled as 1, genes without significant changes were labelled as 0.

