## Supplemental Figures for "CMTR1 is recruited to transcription start sites and promotes ribosomal protein and histone gene expression in embryonic stem cells"

#### Supplemental Figure 1

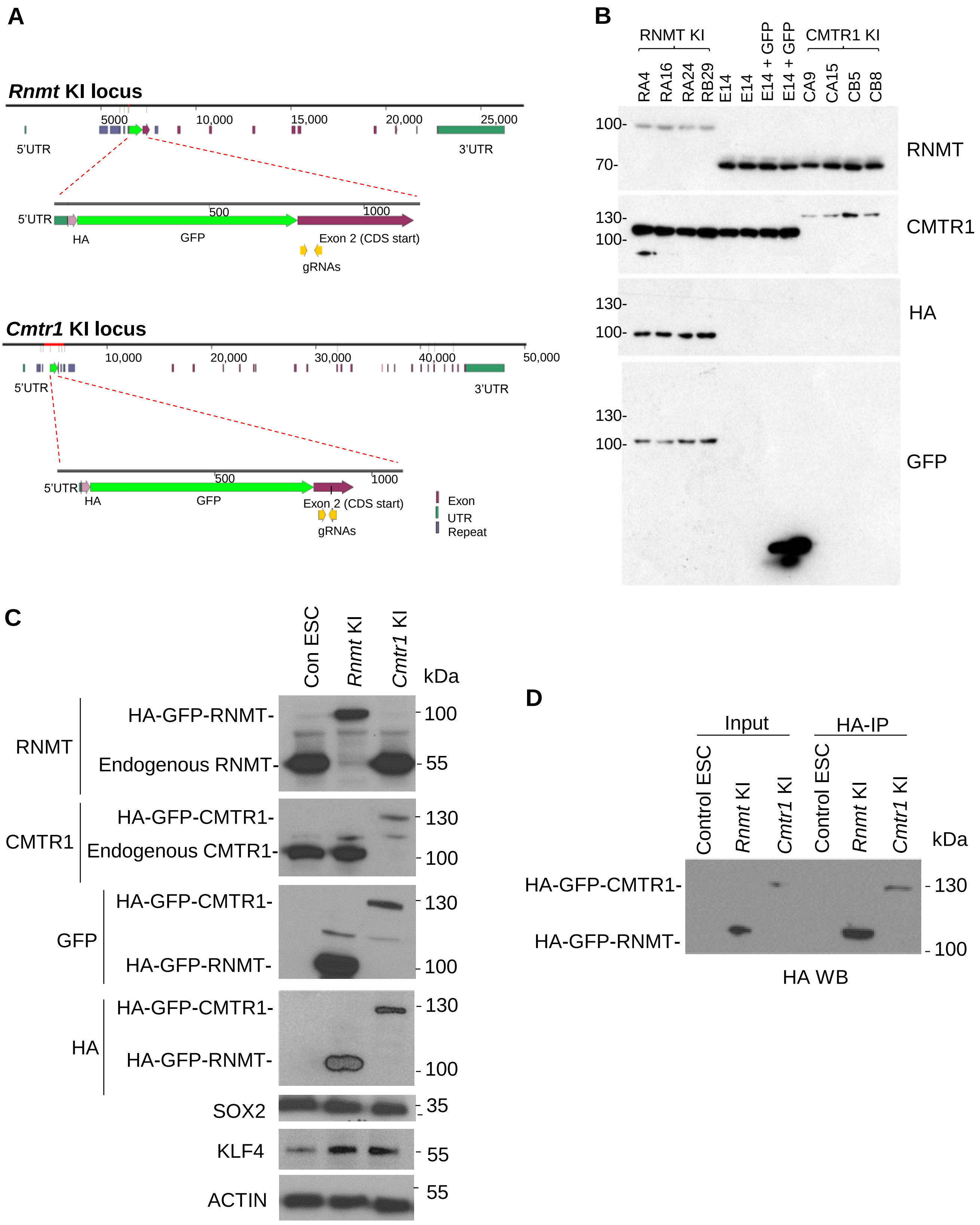


**Figure S1. HA-GFP *Rnmt* and *Cmtr1* KI ESCs** (A) Depiction of the *Rnmt* and *Cmtr1* knock-in (KI) strategy using CRISPR-Cas9. cDNA encoding the HA tag fused to enhanced GFP (HA-GFP) was knocked into the 5' ends of the endogenous *Rnmt* and *Cmtr1* loci. The targeting regions of the guide RNAs (gRNA) and the start of the coding sequence (CDS) are marked. Exons, UTRs (untranslated regions) and Repetitive DNA sequences (Repeats) are depicted. (B)Western blot analysis of 4 clones of each KI and control. (C) Western blot analysis of protein expression in the *Rnmt* and *Cmtr1* HA-GFP KI ESCs. (D) Immunoprecipitation (IP) using anti-HA magnetic beads of HA-GFP-RNMT and HA-GFP-CMTR1 in the *Rnmt* and *Cmtr1* HA-GFP KI ESCs.

### Supplemental Figure 2

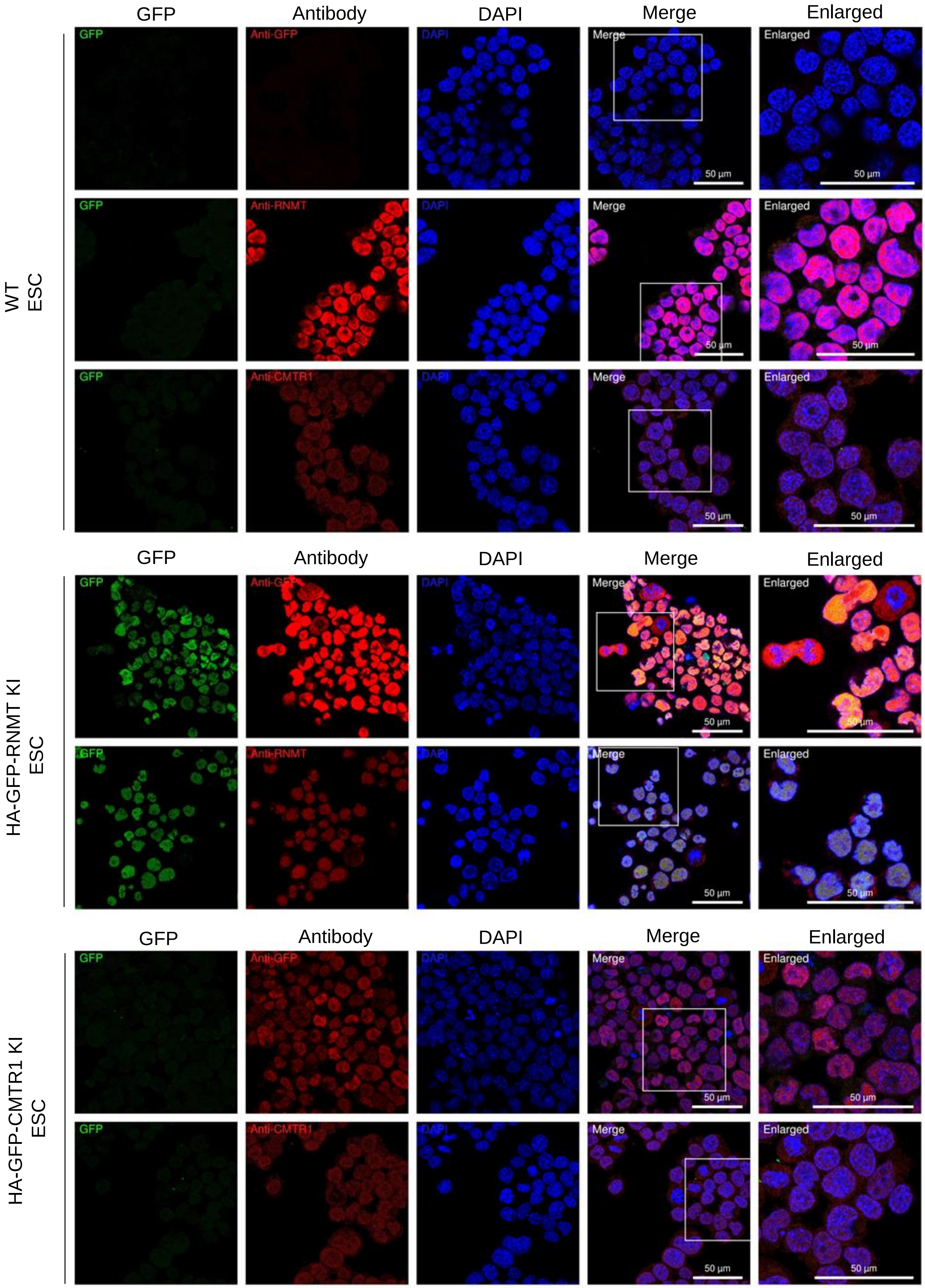


**Figure S2. Endogenous and HA-GFP knock-in RNMT and CMTR1 are predominantly nuclear in ESC lines.** Log phase parental ESC and HA-GFP-RNMT and HA-GFP-CMTR1 lines were fixed and stained with anti RNMT, CMTR1 and GFP antibodies. GFP was also visualised directly.

**A**

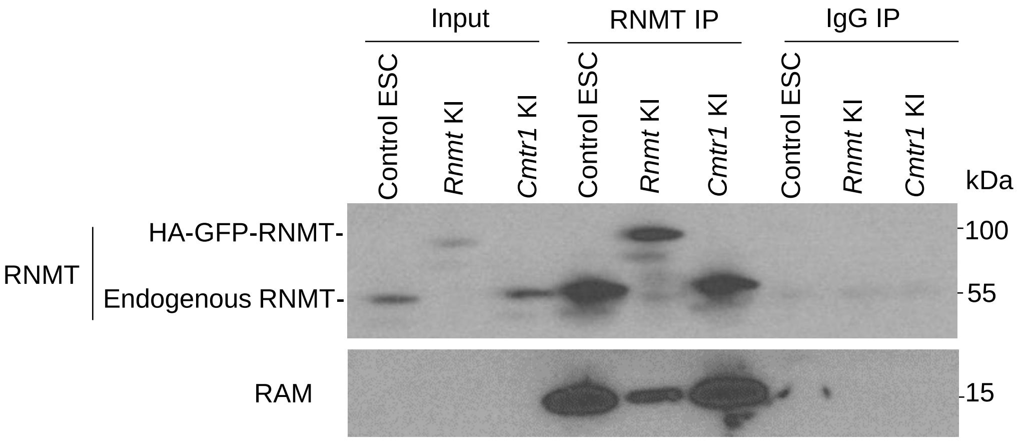


**B**

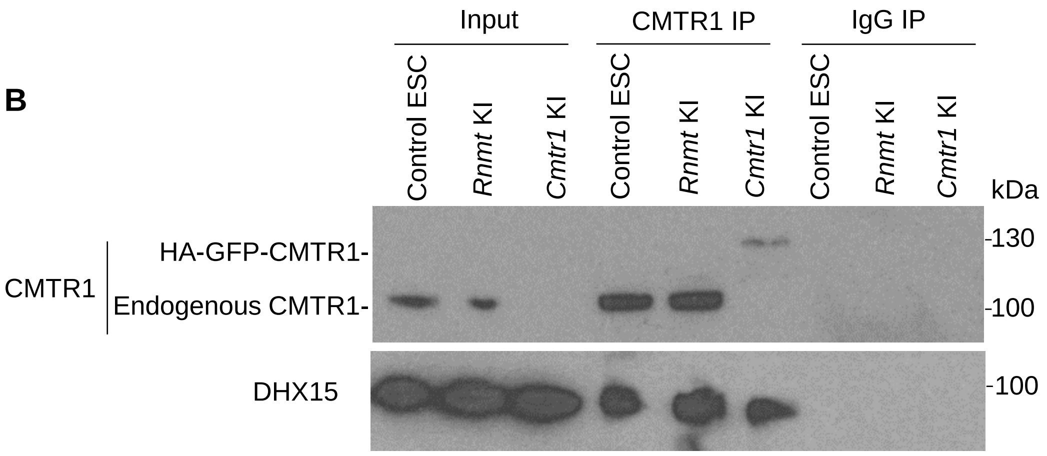


**Figure S3. Endogenous and HA-GFP knock-in RNMT and CMTR1 are co-immunoprecipitated with binding proteins.** Extracts from parental ESC and HA-GFP-RNMT and HA-GFP-CMTR1 knock-in lines were subject to immunoprecipitation using (A) anti-RNMT and (B) anti-CMTR1 antibodies, or IgG antibodies as a control. Western blot analysis was performed

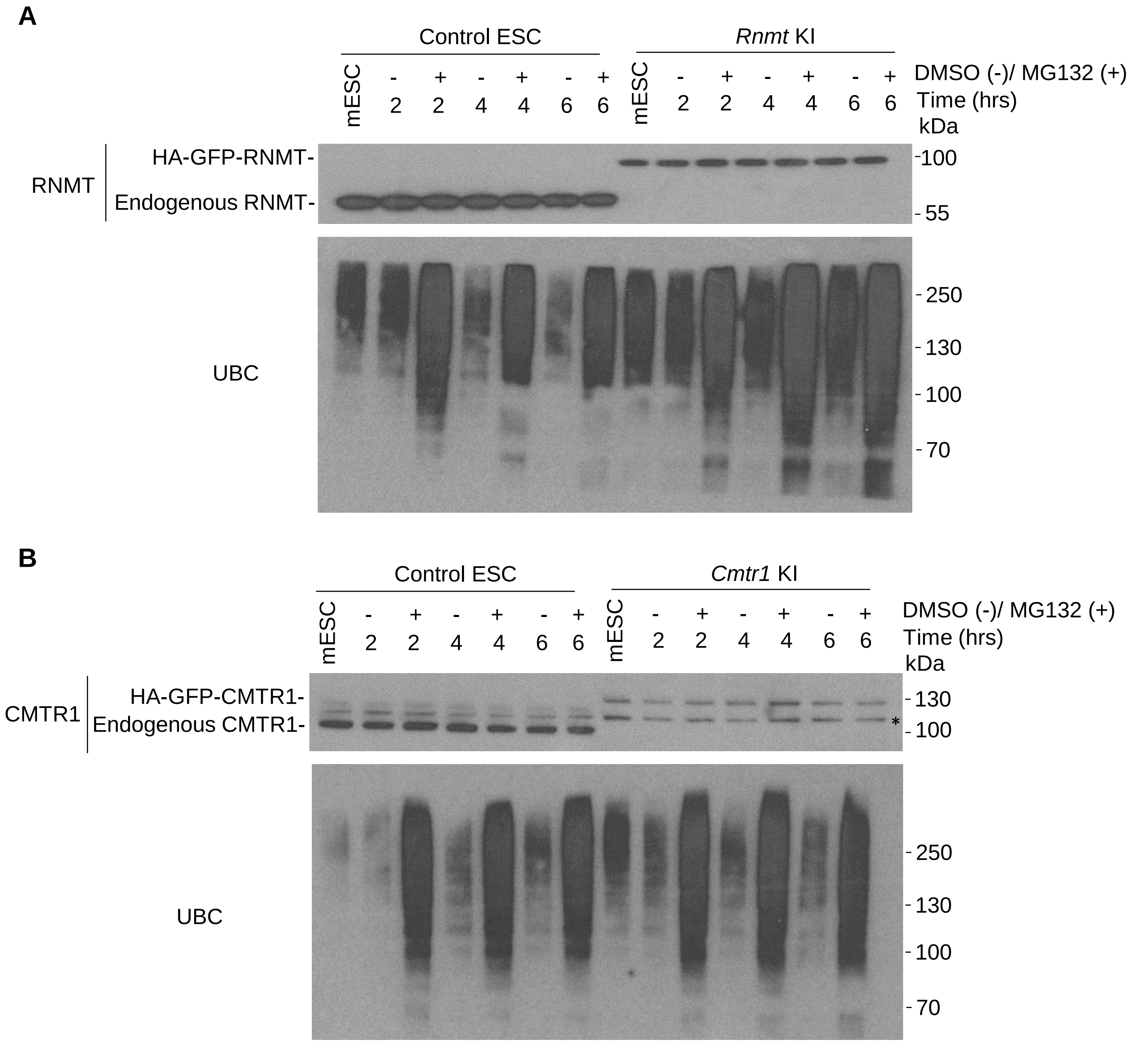


**Figure S4. Endogenous and HA-GFP knock-in RNMT and CMTR1 are not stabilised by incubation with MG132. P**arental ESC and HA-GFP-RNMT and HA-GFP-CMTR1 knock-in lines were treated with 5µM MG132 or DMSO control for the time indicated. Western blots were performed to detect A) RNMT or B) CMTR1. UBC expression was used as a positive control for MG132-dependent stabilisation of proteins. * in CMTR1 indicates non-specific band, verified by CMTR1 IP and KO.

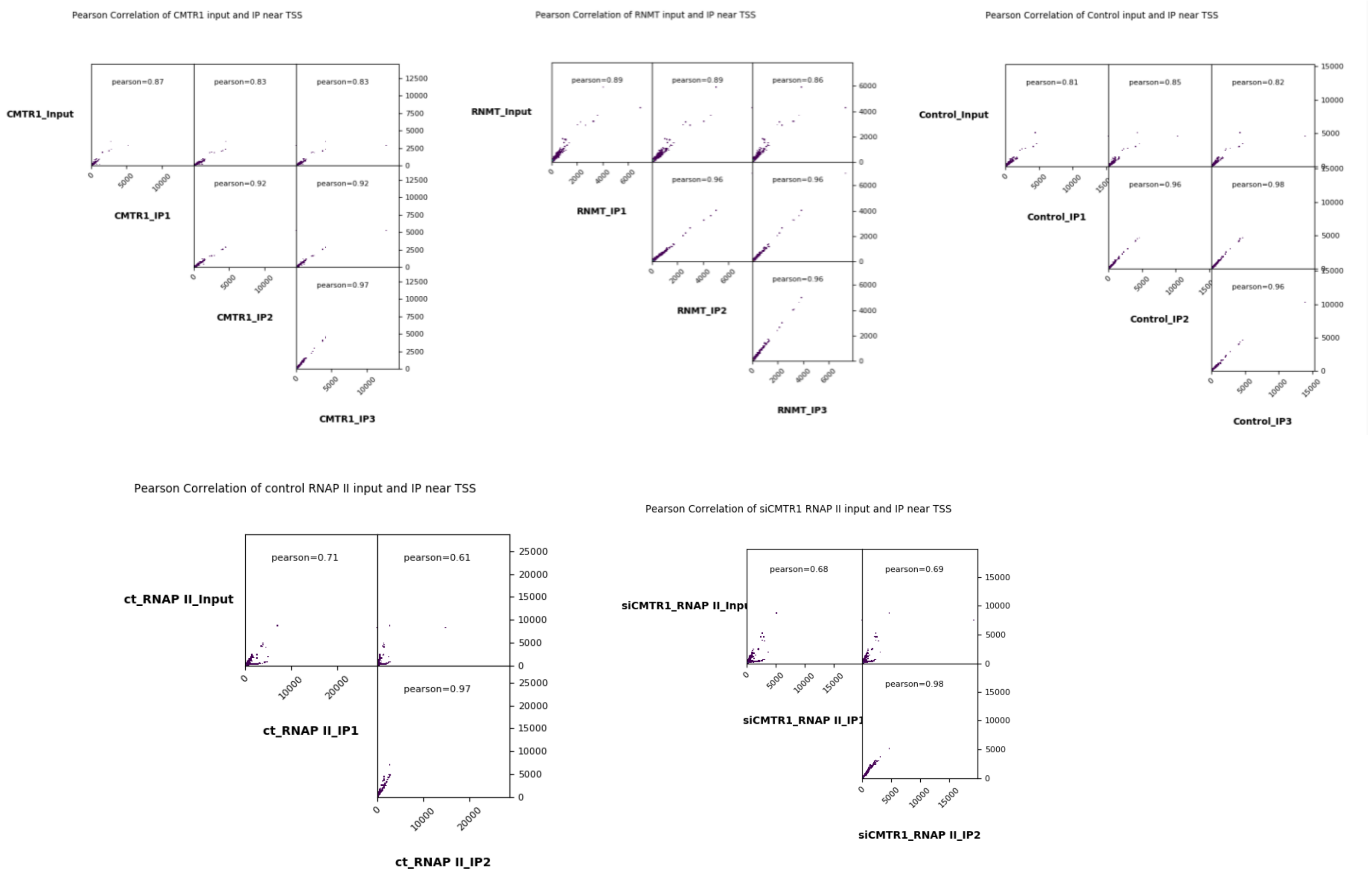


**Figure S5. Pearson correlations of RNA seq and ChIP seq data** Pearson correlation of CMTR1, RNMT, control IP, RNAP II IP (control siRNA) and RNAP II IP (CMTR1 siRNA 1) ChIP-seq library replicates and inputs ±2kb around TSS.

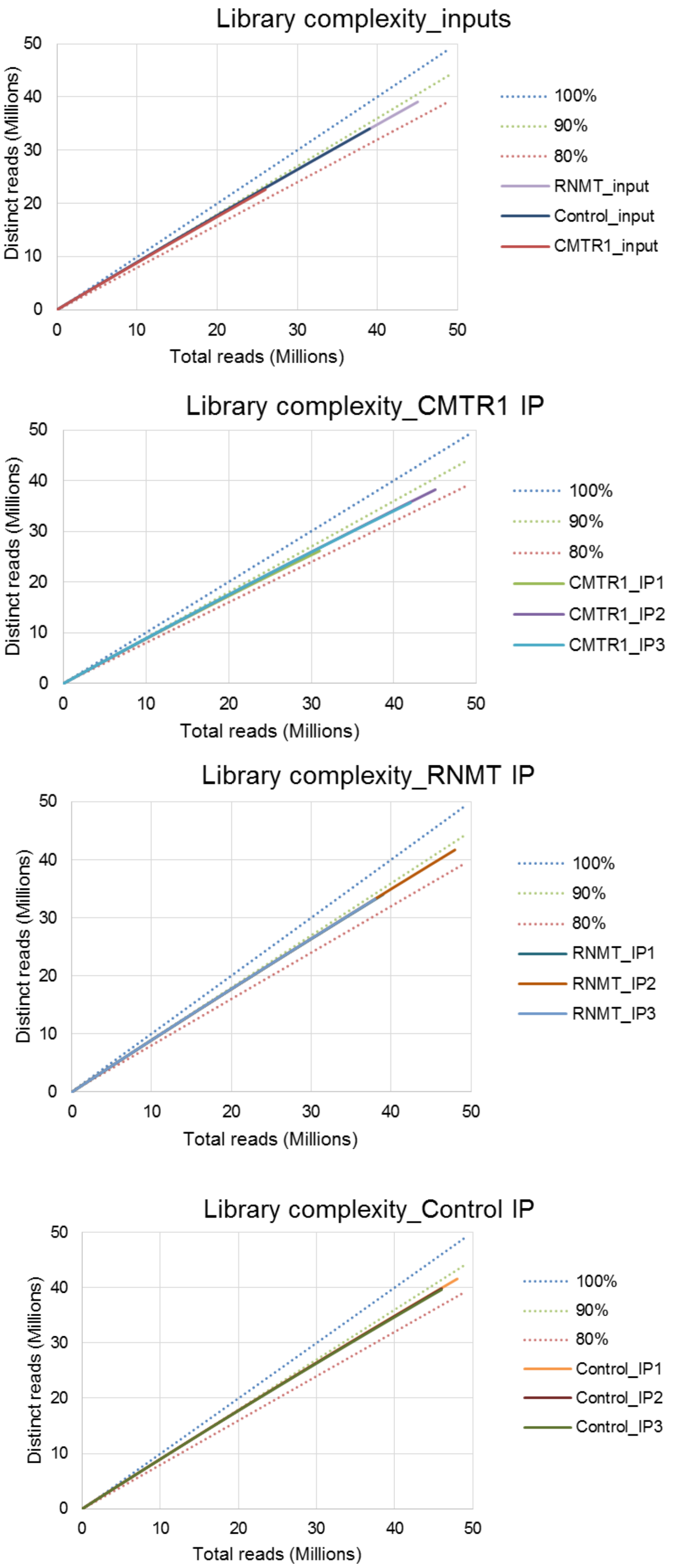


**Figure S6. Line chart indicating the ChIP-seq library complexity** Solid lines indicating the proportion of uniquely mapped reads of all mapped reads. Dash lines marked the boundary of 80%, 90% and 100% uniquely mapped reads.

A

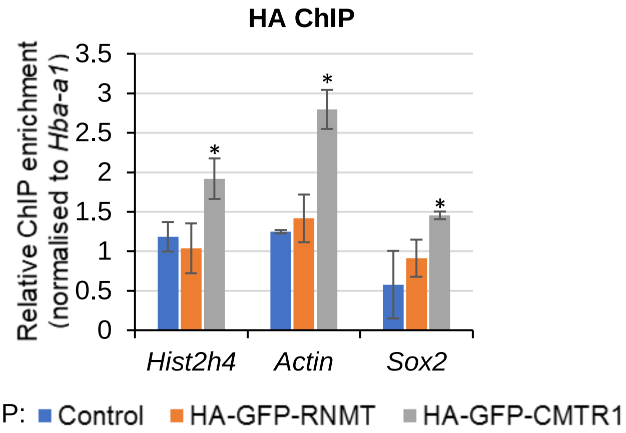


B

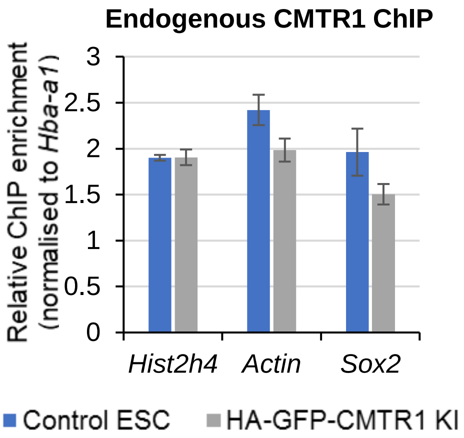


**Figure S7. ChIP-qPCR analysis of HA-GFP-RNMT and HA-GFP-CMTR1 recruitment** (A) ChIP assays were performed using anti-HA-antibody conjugated magnetic beads in ESC control cells, HA-GFP-RNMT KI cells and HA-GFP-CMTR1 KI cells (N=3). T-test P value <0.05 indicated by "*". (B) ChIP assays were performed using the endogenous anti-CMTR1 antibody in ESC control cells and HA-GFP-CMTR1 KI cells (N=2). Relative enrichment of DNA regions bound by the capping enzymes were obtained by calculating the fold change of ChIP signals between the target TSS and the negative control gene *Hba-a1* TSS. Data is presented as average and standard deviation.

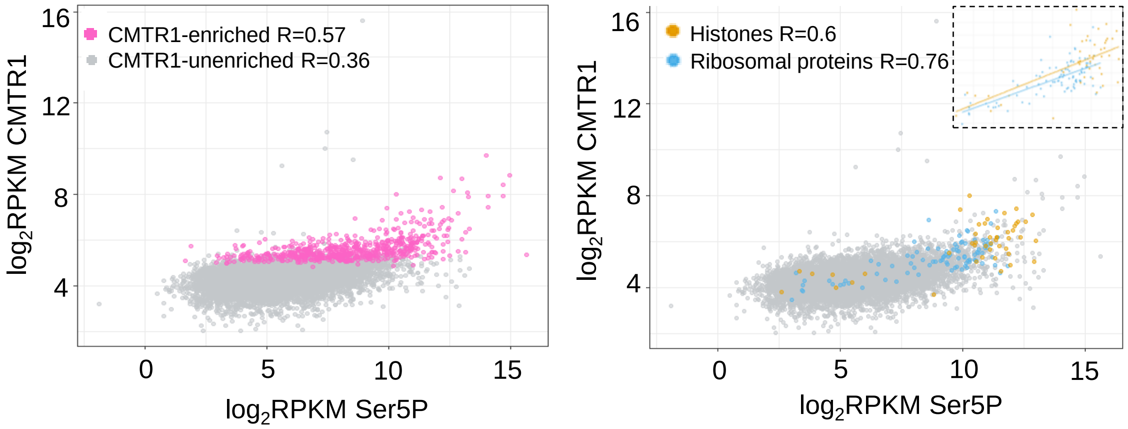


**Figure S8. Correlation of CMTR1 and RNAPII Ser5P ChIP signal** Dot plots indicating the correlation between CMTR1 and RNAPII Ser5P read densities (log$_2$RPKM) ±500b around TSS in the expressed CMTR1-enriched genes (N=768), histones (N=68) and ribosomal proteins (N=100) and all expressed genes in ESC.

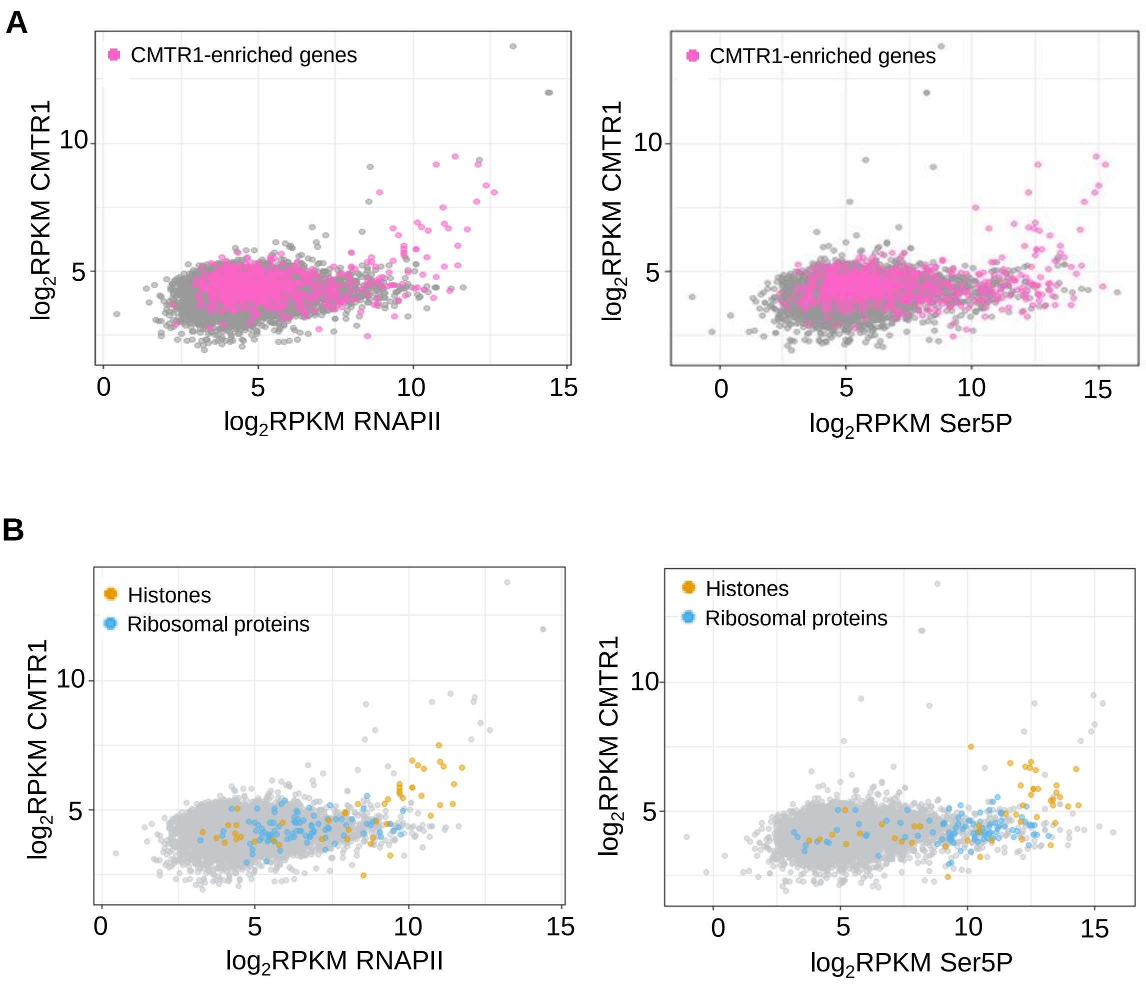


**Figure S9. Correlation of CMTR1 and RNAPII ChIP signal at TES** (A) Dot plots indicating the correlation between CMTR1 and RNAPII (left) or RNAPII Ser5P (right) read densities ($log_2$RPKM) ±500b around TES in the expressed CMTR1-enriched genes (768 genes) and CMTR1-unenriched genes, from all expressed genes in ESC. (B) Dot plots indicating the correlation between CMTR1 and RNAPII (left) or RNAPII Ser5P (right) read densities ($log_2$RPKM) ±500b around TES in histones (68 genes), ribosomal proteins (100 genes) and other expressed genes of all the expressed genes in ESC.

A

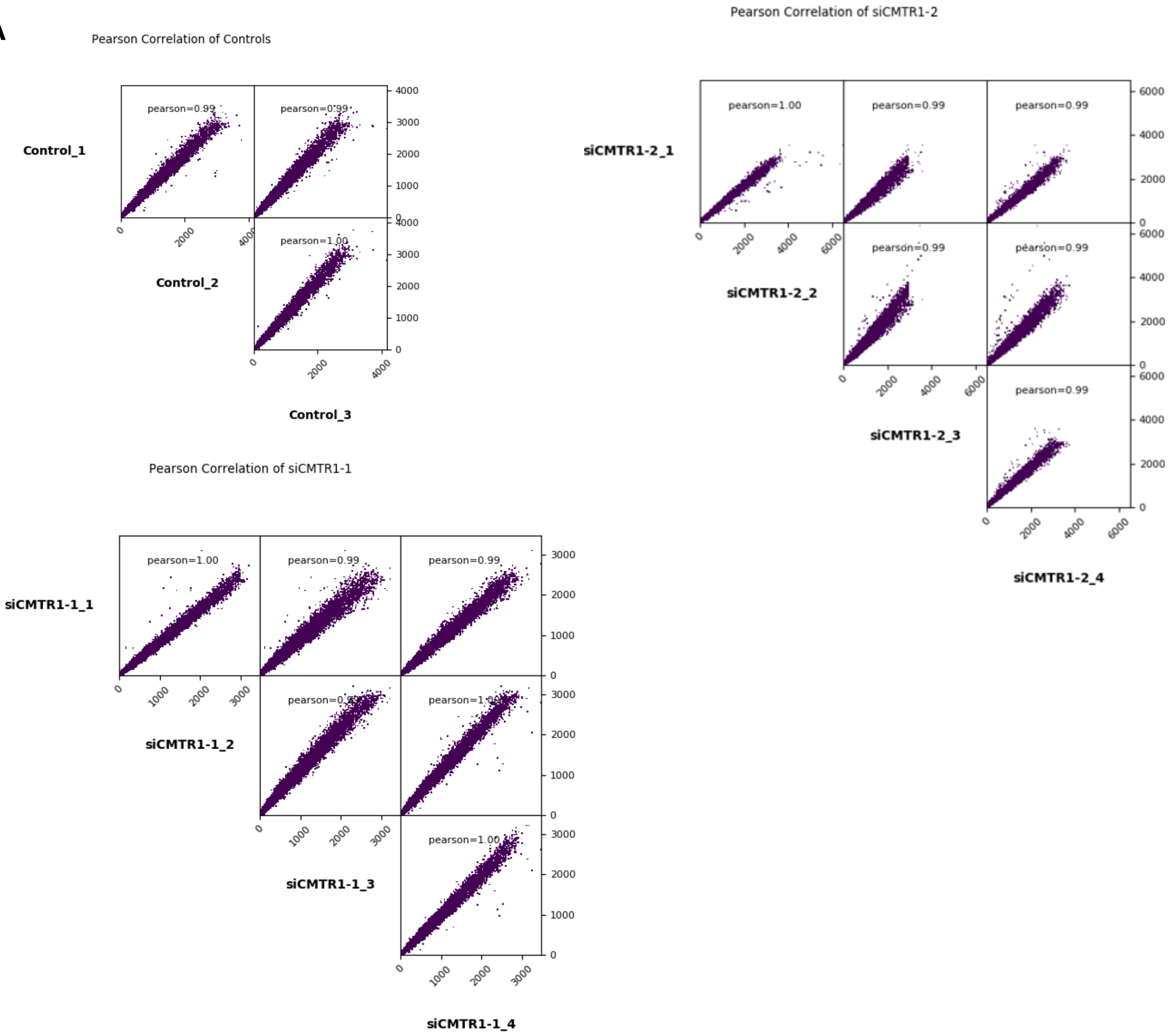


B

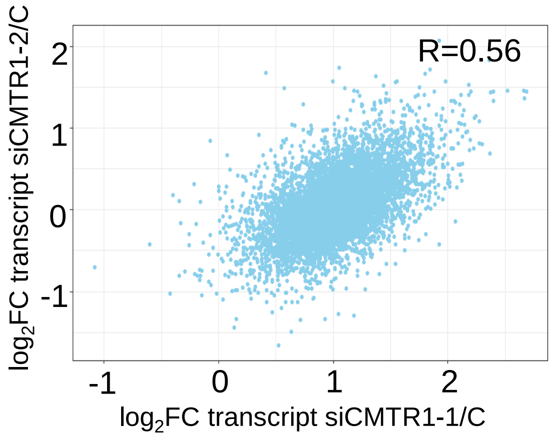


**Figure S10. RNA-seq libraries analysis prepared from ESC transfected with CMTR1 siRNA and control** (A) Pearson correlation of CMTR1 knockdown RNA-seq replicates using the control siRNAs and two siRNAs targeting CMTR1 (1 and 2). (B) Correlation plots of $\log_2$ FC of transcripts between CMTR1 siRNA 1 and 2.

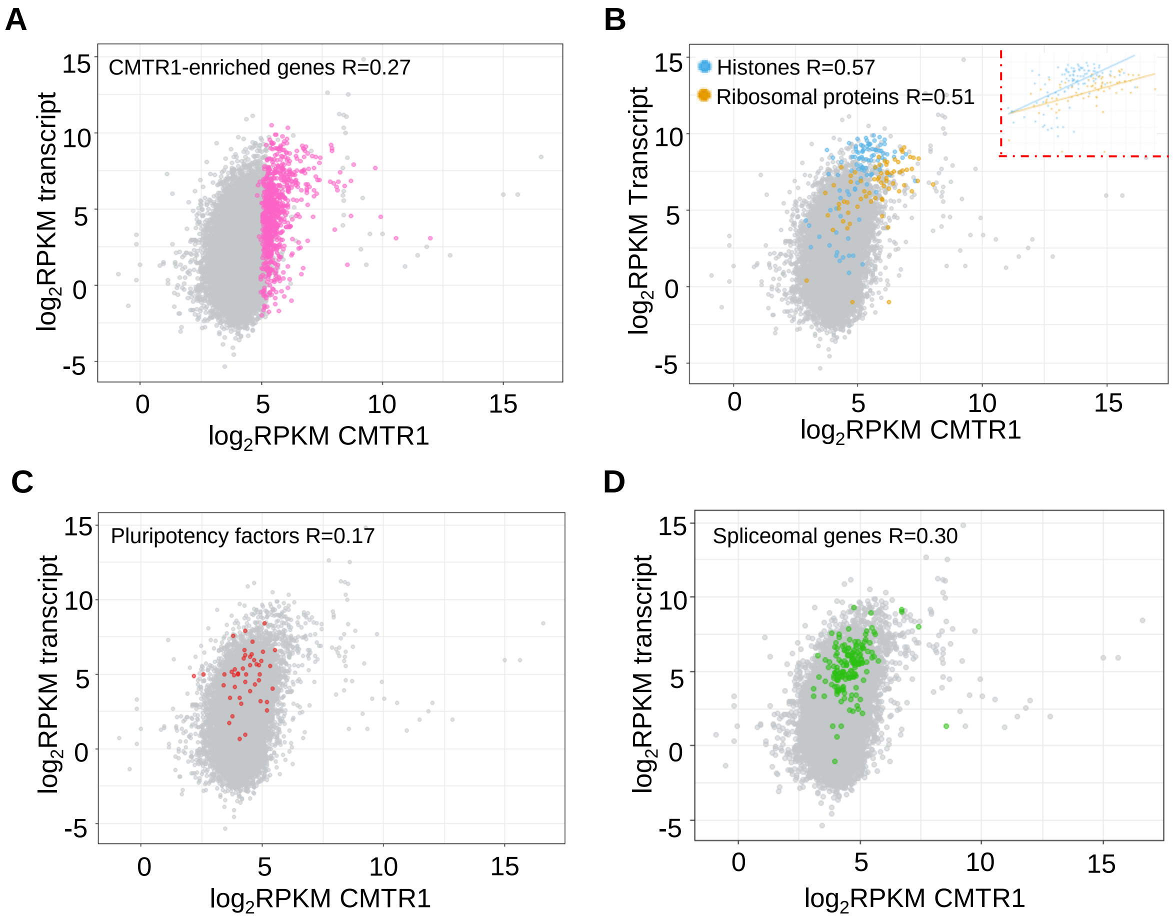


**Figure S11. Correlations between CMTR1 ChIP signal (TSS ±500b) and RNA-seq transcript levels** In CMTR1-enriched genes (768 genes), histones (68 genes) and ribosomal protein genes (100 genes), spliceosomal genes (154 genes) and pluripotency genes (45 genes).

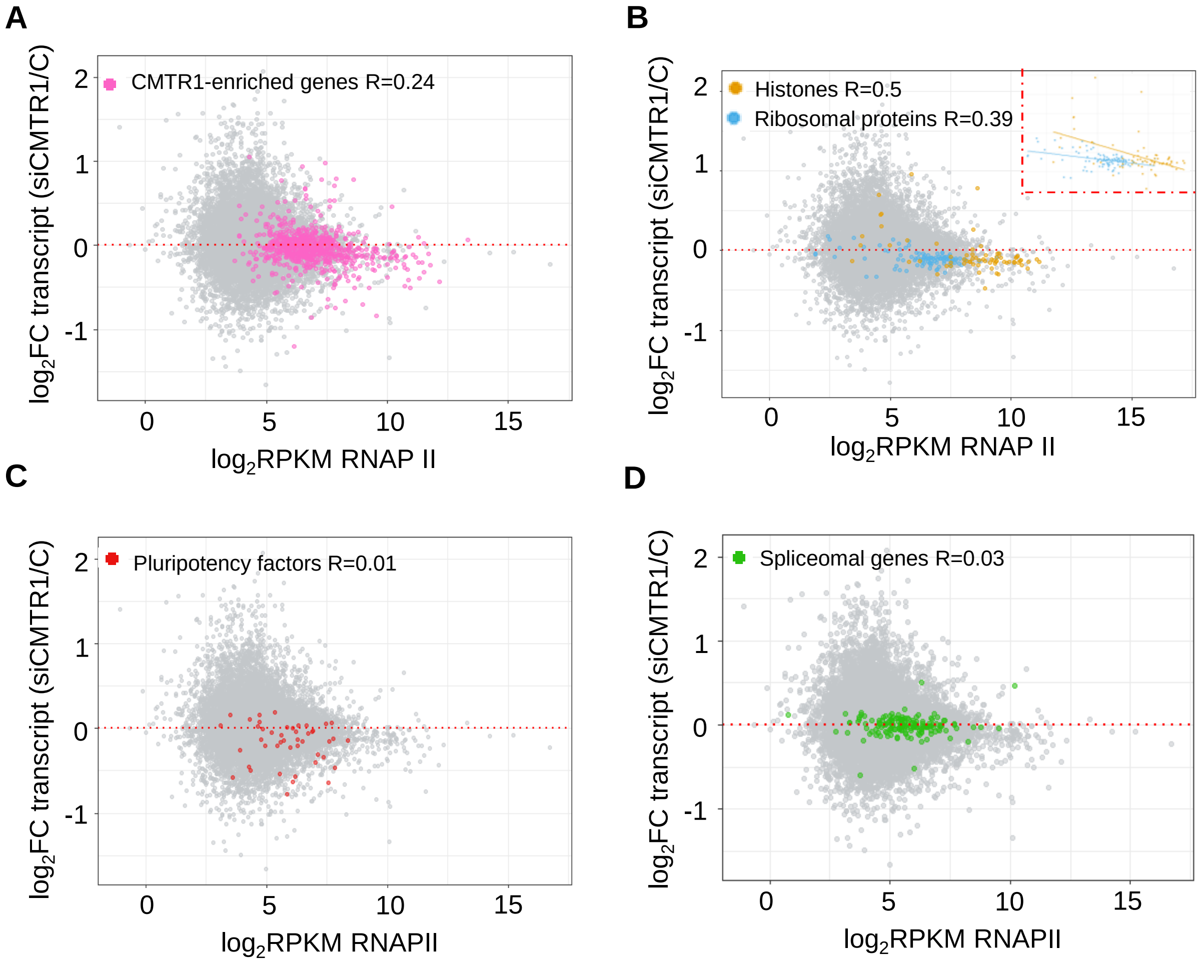


**Figure S12. Correlation of RNAPII levels in control cells and change in transcript level following CMTR1 siRNA transfection** (A-D) Dot plots indicating the correlations between RNAP II bindings (TSS ±500b) and the influence of CMTR1 knockdown on transcript levels in CMTR1-enriched genes (768 genes), histones (68 gene) and ribosomal protein genes (100 genes), spliceomal genes (154 genes) and pluripotency genes (45 genes).

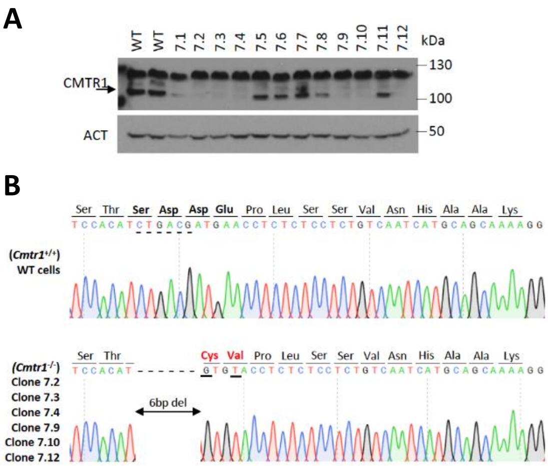


**Figure S13. Analysis of *Cmtr1*$^{KD}$ ESC** CMTR1 protein expression was analysed by western blot in *Cmtr1*$^{KD}$ ESC, (A) from single cell clones. (B) Genomic DNA sequencing analysis.

### Supplemental Figure 14

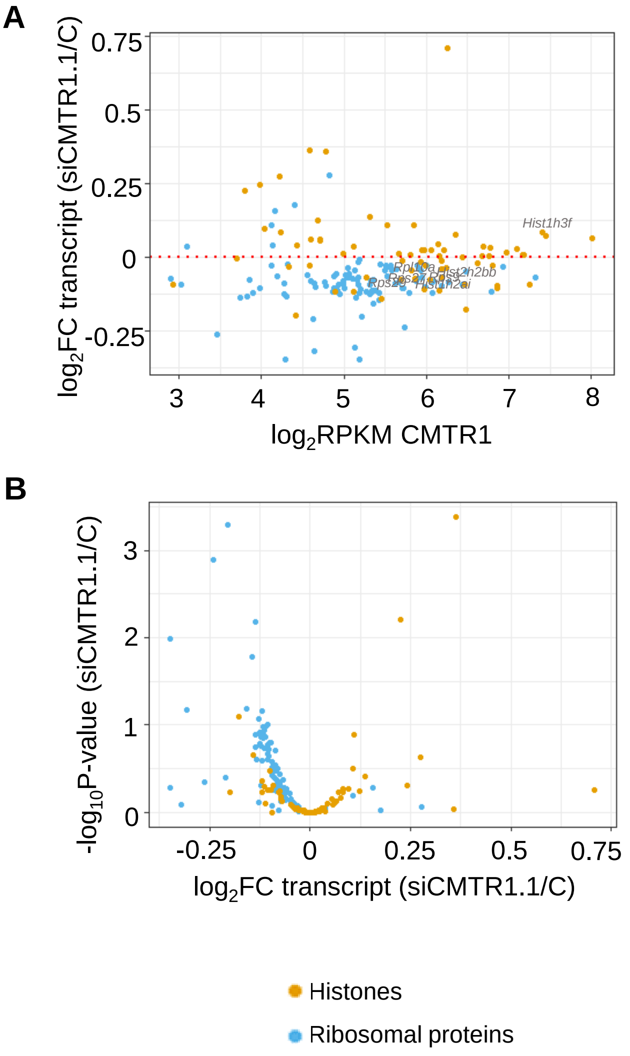


**Figure S14. Histones and ribosomal proteins transcription analysis using different siRNA targeting CMTR1 (CMTR1 si1).** (A) Dot plots indicating gene expression in control cells and Log$_2$FC following CMTR1 siRNA 1 transfection. (B) Volcano plots demonstrating Log$_2$FC expression and EdgeR exactTest –log FDR adjusted P value.

**Supplemental Figure 15**

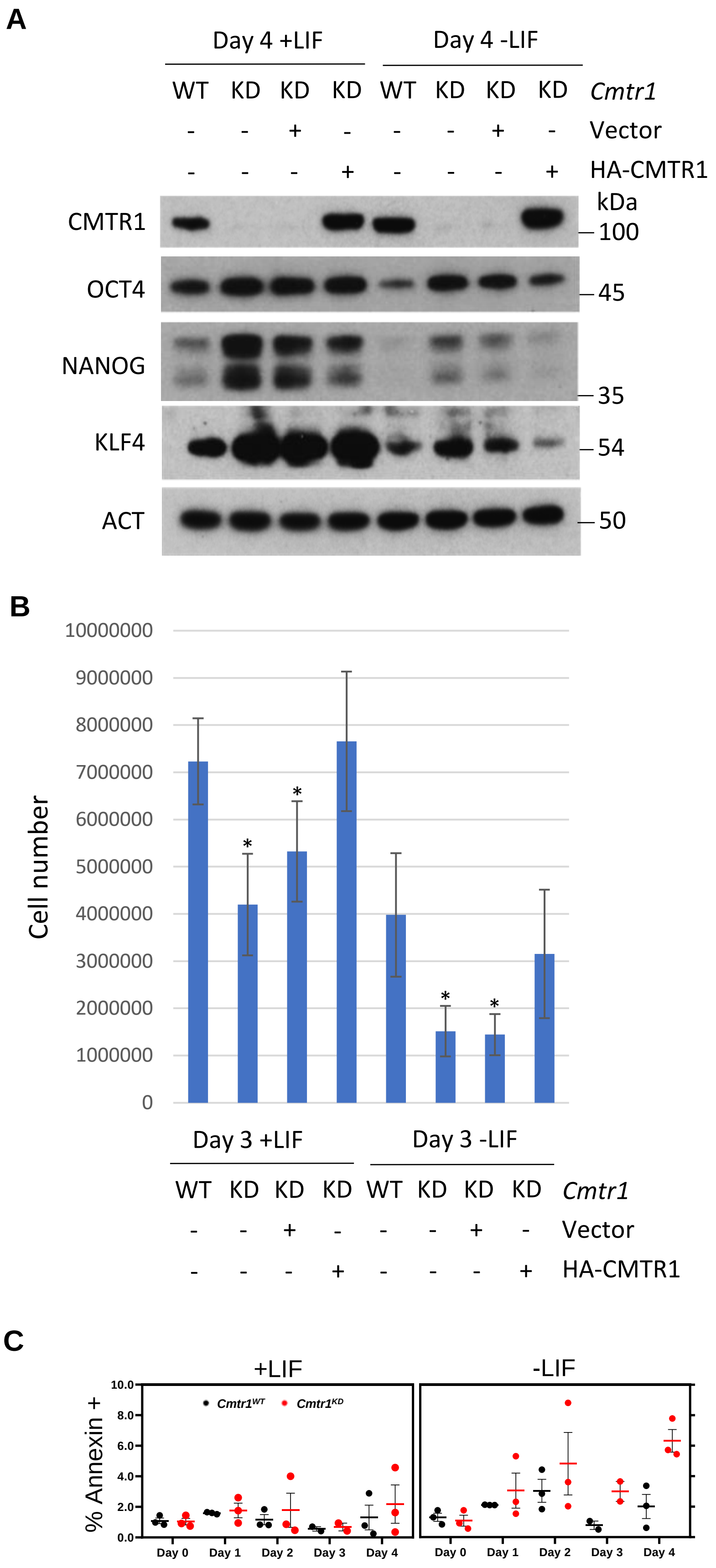


**Figure S15. Rescue of *Cmtr1*$^{KD}$ ESC with HA-CMTR1** *Cmtr1*$^{KD}$ ESC were transfected with pPYPCAGGS-HA-CMTR1 or vector control. ESC (WT) were used as a control. Following LIF withdrawal (-LIF) or control (+LIF) for 3 or 4 days, (A) western blots were performed, (B) cells were counted (n=4). T-test P value<0.05, relative to control cells for that condition, “*”. C) ESC differentiation was induced by LIF withdrawal in *Cmtr1*$^{Wt}$ and *Cmtr1*$^{KD}$ ESCs. Percentage Annexin V positive cells presented (N=3).

#### Supplemental Figure 16

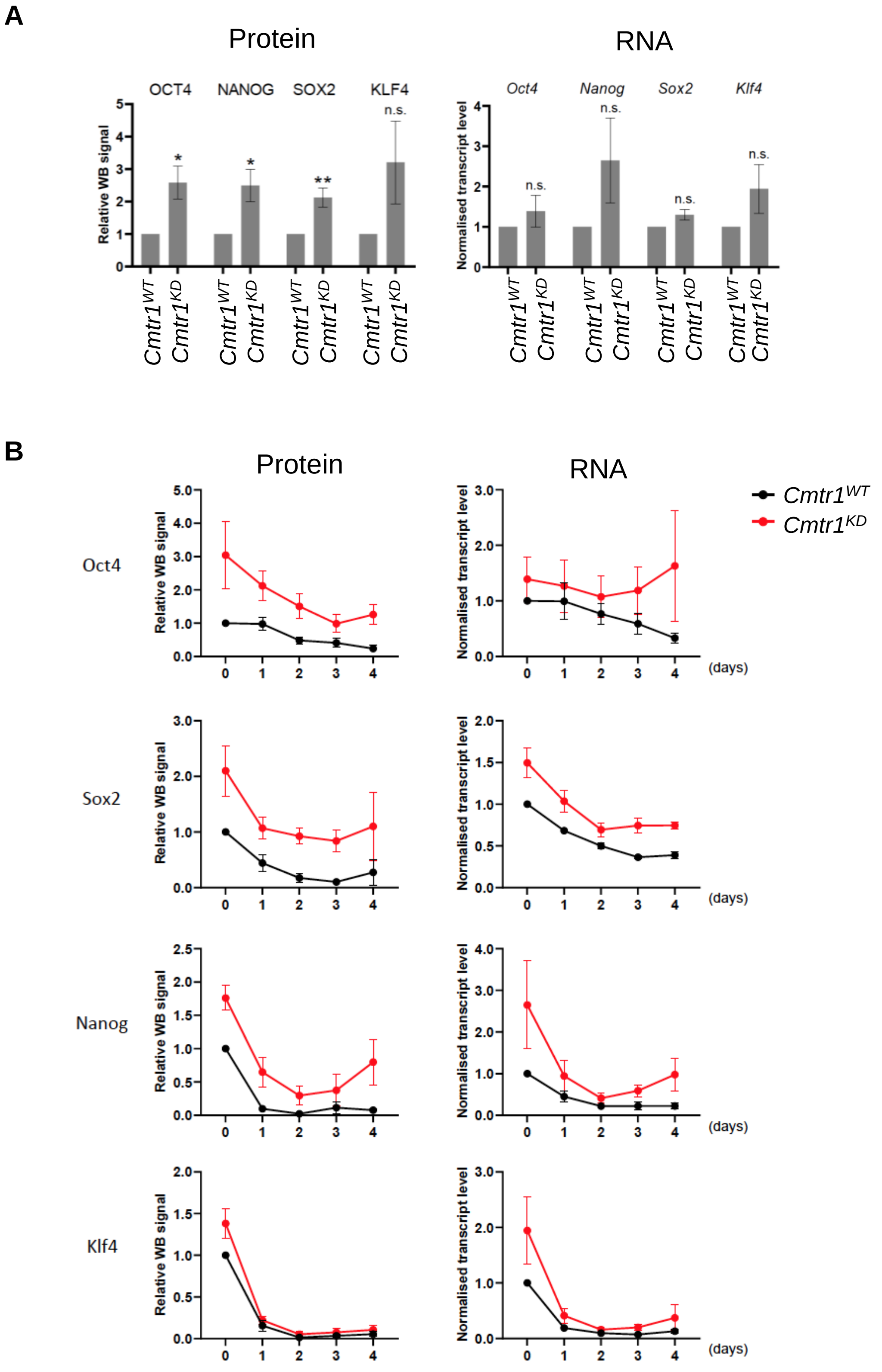


**Figure S16 Protein and RNA levels of pluripotency factors in *Cmtr1*$^{WT}$ and *Cmtr1*$^{KD}$ ESCs**

A) Log-phase *Cmtr1*$^{WT}$ and *Cmtr1*$^{KD}$ ESCs were analysed from protein content by western blot. Levels of proteins indicated were quantified using ImageJ software and normalised to Actin levels. Data is the average and SEM (N=3). RNA levels were quantified by RTPCR and normalised to Actin and Gapdh. Data is the average and SEM (N=4). B) *Cmtr1*$^{WT}$ and *Cmtr1*$^{KD}$ ESCs were cultured in the absence of LIF for 4 days. Protein and RNA analysed as above. Data is average and SEM (N=6)

**A**

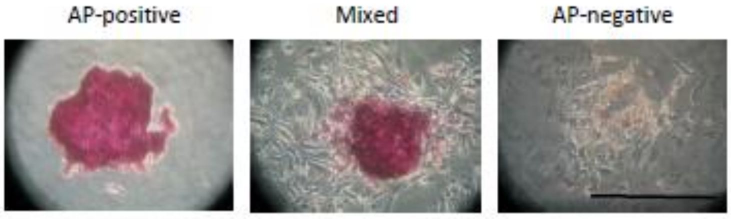


**B**

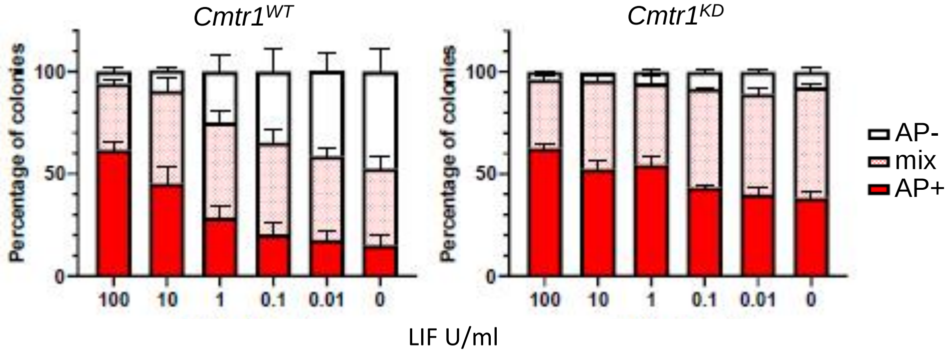


**Figure S17 Impact of LIF withdrawal on *Cmtr1$^{Wt}$* and *Cmtr1$^{KD}$* ESCs.**
A,B) ESC were plated on gelatinised 6-well plates. After 3 days, cells were cultures in a titration of 0-100 units/ml LIF. After 6 days cells were fixed and stained for alkaline phosphatase. (A) Colonies were stained as alkaline phosphatase positive, negative or mixed. Examples provided. B) Chart of percentage of each colony type. Data presented is average and SEM (N=3).
